# High-resolution QTL mapping with Diversity Outbred mice identifies genetic variants that impact gut microbiome composition

**DOI:** 10.1101/722744

**Authors:** Florencia Schlamp, David Y Zhang, Juan Felipe Beltrán, Elissa J Cosgrove, Petr Simecek, Matthew Edwards, Julia K Goodrich, Ruth E Ley, Allan Pack, Gary A Churchill, Andrew G Clark

## Abstract

The composition of the gut microbiome is impacted by a complex array of factors, from nutrient composition and availability, to physical factors like temperature, pH, and flow rate, as well as interactions among the members of the microbial community. Many of these factors are affected by the host, raising the question of how host genetic variation impacts microbiome composition. Though human studies confirm this type of role for host genetics, its overall importance is still a subject of debate and remains difficult to study. The mouse model, by allowing the strict control of genetics, nutrition, and other environmental factors, has provided an excellent opportunity to extend this work, and the Diversity Outbred (DO) mice in particular present a chance to pinpoint host genetic variants that influence microbiome composition at different levels of generality. Here, we apply 16S rRNA gene sequencing to fecal samples of 247 DO male mice to estimate heritability and perform taxon-specific QTL mapping of microbial relative abundances revealing an increasingly heterogeneous picture of host function and microbial taxa at the host-microbiome interface. We present the first report of significant heritability of phylum Tenericutes in mice, and find novel QTL-spanning genes involved in antibacterial pathways, immune and inflammatory disease, and lipid metabolism.

## INTRODUCTION

The gastrointestinal tract of all vertebrates, including humans, harbors a complex ecological community of highly diverse microbes referred to as the gut microbiota. The microbiota colonizes the gut for the first time during the birth of the host, and its composition is influenced by many factors during the host’s life such as disease, diet, and antibiotics (Francino 2016; Battaglioli AND Kashyap 2018; Dudek-Wicher *et al.* 2018; Dash *et al.* 2019). Variation in the human gut microbiome composition has also already been associated with host immune responses (Round AND Mazmanian 2009; Garrett *et al.* 2010; Veiga *et al.* 2010), metabolic phenotypes (Turnbaugh *et al.* 2009; Ridaura *et al.* 2013), and diseases such as obesity (Ley *et al.* 2005), heart disease (Fava *et al.* 2006), and diabetes (Wen *et al.* 2008). Given the roles of the gut microbiome in complex human diseases, it is important to characterize the factors that impact microbiome composition.

While it is clear that the gut microbiome composition is strongly impacted by environmental exposures (Rothschild *et al.* 2018), the role of host genetics has only recently been implicated (Goodrich *et al.* 2014; Blekhman *et al.* 2015; Goodrich *et al.* 2016). Studies have identified multiple genetic variants significantly associated with specific bacterial taxon abundances (Davenport *et al.* 2015; Bonder *et al.* 2016; Turpin *et al.* 2016; Wang *et al.* 2016; Goodrich *et al.* 2017; Igartua *et al.* 2017; Rothschild *et al.* 2018) despite the observation that generally the primary determinants of microbiome composition are non-genetic (Rothschild *et al.* 2018). The relationship between genetic and non-genetic determinants is complex, as in the case of diet, which can influence the variability of complex traits by reshaping the gut microbiome (Vorobyev *et al.* 2019). Overall, it is clear that host-microbiome relationships are impacted by interactions between genetics and environment to drive both community composition and host traits (Kurilshikov *et al.* 2020). Human genetic studies have significant limitations for accurate assessment of genetic effects on the microbiome, including accessibility to large and diverse sample populations as well as a general lack of control over confounding variables like diet, thus only detecting the strongest genetic effects. This lack of experimental control can be circumvented through studies in model organisms, which would allow us to better characterize host-microbiome relationships and increase our chances of identifying genetic effects.

The mouse model, with the ability to control diet, provides a better opportunity to dissect genetic and environmental factors impacting microbiome composition and has been successful in this endeavor using inbred strains. Quantitative trait locus (QTL) mapping efforts show that gut microbiota composition is a polygenic trait, with clearly mappable genetic factors influencing the gut microbiome composition (Benson *et al.* 2010; Mcknite *et al.* 2012; Snijders *et al.* 2016). Standard QTL mapping approaches have low mapping resolution, however, and advanced intercross lines provide one excellent means of improving mapping resolution. Belheouane *et al.* (2017) performed genetic and 16S rRNA gene analysis of skin microbiomes of a collection of 15-generation advanced intercross lines, and demonstrated that the improved mapping resolution also improved the specificity and significance of genetic associations. It is clear that the mouse model will provide further opportunities to dissect the means by which the host genome can modulate microbiome composition. A logical next step is a mapping experiment to identify portions of the genome that influence functional pathways that modulate the microbiome.

Here we extend the analysis of the link between the host genome and microbiome using the Diversity Outbred mouse model. The Diversity Outbred (DO) population is a heterogeneous mouse stock derived from the same eight progenitor lines (A/J, C57BL/6J, 129S1/SvImJ, NOD/ShiLtJ, NZO/HlLtJ, CAST/EiJ, PWK/PhJ, and WSB/EiJ) used to establish the Collaborative Cross (CC) (Collaborative Cross Consortium 2012). Mice from the CC lines at early stages of inbreeding were used to establish the DO population, which is maintained by randomized outbreeding among 175 mating pairs. The result is each individual DO mouse represents a unique combination of segregating alleles drawn from the original eight progenitor lines. The advantages of this outbreeding include normal levels of heterozygosity — similar to the human genetic condition — and substantially increased genetic resolution (Churchill *et al.* 2012). Both the DO mice and their founder progenitor lines have already proven to be successful in identifying genetic associations with intestinal microbiome composition (O’Connor *et al.* 2014, Kemis *et al.* 2019).

In this study, motivated by the high level of environmental control of the laboratory mouse and the improved mapping resolution of the Diversity Outbred mouse system, we identified genetic underpinnings of the gut microbiota of 247 Diversity Outbred mice. We uncover evidence of host genetic factors influencing the composition of many specific attributes of the gut microbiome (**Figure 1**). These included not only associations between specific host genetic variants and abundances of particular bacterial taxa, but also associations with functional molecular pathways.

**Figure 1.**
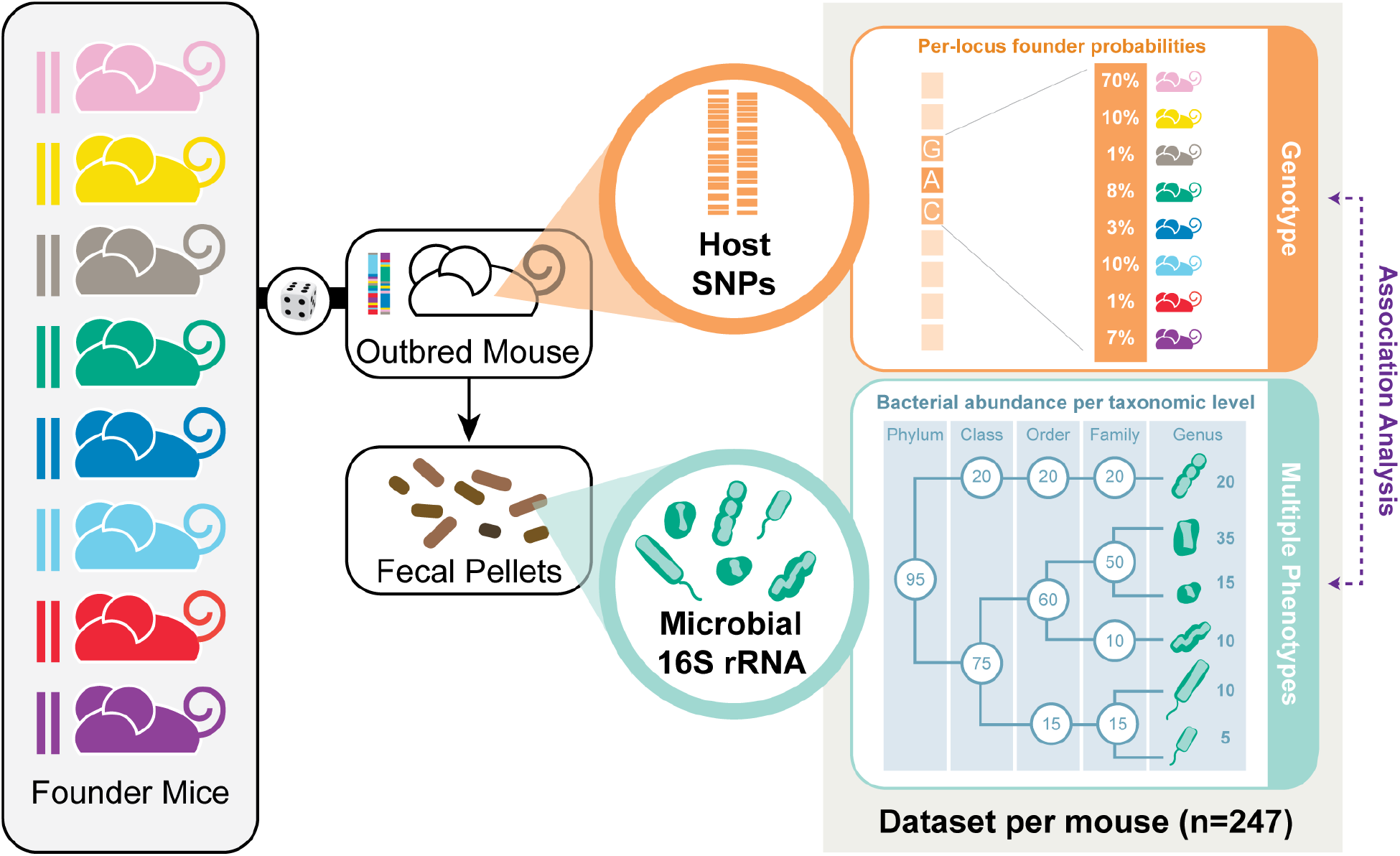
Data flow schematic. Each of the 247 mice in this study represents a unique combination of segregating alleles, whose genome is a unique sampling from the original eight progenitor lines (founder mice). The SNP genotype of each mouse is represented by an eight-founder state probability. Microbial 16S rRNA from fecal pellets from each DO mouse provided bacterial relative abundances, which were aggregated at each taxonomic level and used as separate phenotypes/traits.

## MATERIALS AND METHODS

### Animal population and sample collection

Male mice from the Diversity Outbred Mouse Panel were obtained from The Jackson Laboratory (Bar Harbor, ME, USA) at 6 weeks of age. Experiments were performed at the University of Pennsylvania, Center for Sleep. Mice were group-housed (5 animals per cage) for 2 weeks of post-travel acclimation, and then single-housed at identical conditions with lights on/lights off at 7:00 AM/7:00 PM with a lux level of 60 and temperature 23-25°C. Bedding used in home cages was Bed-o Cobs 1/8” (The Andersons Inc., Maumee, OH). Mice were fed *ad libitum* Laboratory Autoclavable Rodent Diet 5010 (Lab Diet, St. Louis, MO). Fecal pellets from 249 mice were collected at 3 months old (two samples were later discarded, leaving a final analyzed dataset of 247 mice). The pellets were collected from the mouse cage at 10:00 AM, i.e., 3 hours after lights on. Pellets were stored in Eppendorf tubes placed on dry ice and moved to a −80°C freezer until shipping and processing at Cornell University (Ithaca, NY, USA).

### Microbial DNA extraction, 16S rRNA gene PCR, and sequencing

Microbial community DNA was extracted from one single frozen pellet per sample using the MO BIO PowerSoil-htp DNA Isolation Kit (MO BIO Laboratories, Inc., cat # 12955-4), but instead of vortexing, samples were placed in a BioSpec 1001 Mini-Beadbeater-96 for 2 minutes. We used 10-50 ng of sample DNA in duplicate 50 μl PCR reactions with 5 PRIME HotMasterMix and 0.1 μM forward and reverse primers. We amplified the V4 region of 16S rRNA gene using the universal primers 515F and barcoded 806R and the PCR program previously described Caporaso *et al.* (2011), but with 25 cycles. We purified amplicons using the Mag-Bind® E-Z Pure Kit (Omega Bio-tek, cat # M1380) and quantified with Invitrogen Quant-iT™ PicoGreen® dsDNA Reagent, and 100 ng of amplicons from each sample were pooled and paired end sequenced (2×250bp) in two separate sequencing runs on an Illumina MiSeq instrument at Cornell Biotechnology Resource Center Genomics Facility.

### 16S data processing

We performed demultiplexing of the 16S rRNA gene sequences and OTU picking using the open source software package Quantitative Insights Into Microbial Ecology (QIIME) version 1.9.0 with default methods (Caporaso *et al.* 2010). The total number of sequencing reads was 15,149,384, with an average of 61,334 sequences per sample and ranging from 17,658 to 135,803. Open-reference OTU picking at 97% identity was performed against the Greengenes 8_13 database. 12% of sequences failed to map in the first step of closed-reference OTU picking. The taxonomic assignment of the reference sequence was used as the taxonomy for each OTU. ‘NR’ within taxa names represents New Reference OTUs defined as those with sequences that failed to match the reference and are clustered *de novo*. Random subsamples were used to create a new reference OTU collection and ‘NCR’ represents New Clean-up Reference OTUs that failed to match the new reference OTU collection (Rideout *et al.* 2014).

For the non-rarefied data, read count was used as an additional covariate during QTL mapping to reduce the effect of sequencing depth. A rarefied dataset was also used for heritability estimates and QTL mapping, as explained in **File S1**. Two extreme outliers were omitted from further analysis, yielding a total of 247 samples. To differentiate the non-rarefied taxa from the rarefied taxa, we use ‘NonR’ to represent the non-rarefied dataset and ‘R’ to represent the rarefied dataset.

For heritability estimates and QTL mapping, a filter was applied across all 247 samples that removed any taxon that was not present in more than 50% of the samples. Relative abundance of reads (number of reads clustered to each taxa divided by the total number of reads in a given sample) was used as the tested phenotype. Relative abundances were rank Z-score transformed using R-package *DOQTL* (Gatti *et al.* 2014).

Stacked bar plots of the most abundant taxa within each taxonomic level were plotted with R-package *ggplot2*. A box-plot was first generated for each taxonomic level depicting the relative abundances of the taxa within that taxonomic level across the 247 samples (**Figure S1**). The top ten taxa with the highest average relative abundances are selected to be plotted in the stacked bar plot, ordered by the most abundant taxon. A heatmap that correlates similarities between taxa from the non-rarefied and rarefied datasets based on the Pearson correlation coefficient was plotted using the R-package *corrplot* (**Figure S3**).

### SNP genotyping

SNP genotyping was done at the Jackson Laboratories on each of the 247 mice using The Mega Mouse Universal Genotyping Array (MegaMUGA). A total of 57,973 SNPs passed QC metrics and were used in the heritability and mapping analysis reported here.

### Heritability estimation

Heritabilities of the various bacterial taxa were quantified and calculated on autosomes using a linear mixed model as implemented in R-package *lme4qtl* via the relmatLmer() function (Ziyatdinov *et al.* 2018) (https://github.com/variani/lme4qtl). This linear mixed model enables us to decompose variability into genetic and environmental components. The variance of the genetic component is expected to be 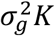, where *K* is a kinship matrix normalized as proposed in (Kang *et al.* 2010). The kinship matrix is specified via the “relmat” argument in relmatLmer(). To account for the potentially confounding effects of shared cages during acclimation (as noted above under **Animal population and sample collection**), we also included cage as a random effect in our model. Thus, the model included estimates of variance of the genetic component 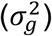 and the cage component 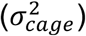, and the residual variance due to unspecified environmental factors 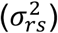. The narrow sense heritability was then estimated as:

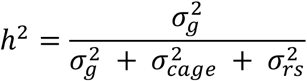

Sequencing run was included as a covariate in both non-rarefied and rarefied datasets. For our non-rarefied dataset, narrow sense heritabilities were calculated using the number of read counts as an additional covariate. Significance of heritability estimates was assessed by conducting a restricted likelihood ratio test using the exactRLRT() function in the R-package *RLRsim* (Scheipl *et al.* 2008), as applied in Supplementary Note 3 in Ziyatdinov *et al.* (2018). We calculated standard errors for the heritability estimates following code posted on the *lme4qtl* GitHub page: https://github.com/variani/lme4qtl/blob/master/demo/se.R. This script uses the deltamethod() function in the R-package *msm* (https://github.com/chjackson/msm) to approximate standard errors using the delta method. Proportion variance estimates for kinship and cage for all taxa and their taxonomic level for rarefied data are presented in **Figure S4**. A comparison of heritability estimates and standard error between non-rarefied and rarefied data can be seen in **Figure S5**.

### QTL Mapping

For QTL mapping, rank Z-score transformed relative abundances were mapped using a linear mixed model in R-package *lme4qtl::relmatLmer()* (Ziyatdinov *et al.* 2018) (fit using maximum likelihood (ML), REML=F) on autosomes with kinship included as a random effect to account for genetic relatedness among animals. For the bacterial taxa from the five taxonomic levels, we generated QTL mappings with the taxa designated as the phenotype. Sequencing run (fixed effect) and cage (random effect) were included in both non-rarefied and rarefied datasets. We included read count as an additional covariate (fixed effect) for our non-rarefied dataset. Significant and suggestive associations were identified in a two-step procedure. First, we applied likelihood ratio tests comparing models with and without genotype. *P*-values derived from these tests were adjusted for multiple testing across SNPs (within a given taxon) using R function p.adjust() with method “BH” (Benjamini AND Hochberg 1995). In the second step, we conducted permutation tests (1000 permutations) for taxa that had associations with adjusted *p*-value < 0.1 in the maximum likelihood analysis. Due to the computational cost of performing permutation tests for each taxa/peak combination, we further filtered the permutation candidates by only querying the peak with the lowest likelihood p-value in regions with peak overlaps. This resulted in permutation *p*-values for 4 taxa and 4 peaks (**Table 2**). Annotated genes found within QTL regions with permutation *p*-value < 0.1 can be found in **Table S5**. Although *p*-values are corrected within each trait, no additional adjustment is made for the search across traits.

**Table 2.** QTL regions for OTUs. Only showing OTUs with adj. *p*-value < 0.1 (statistically suggestive) and with a QTL region overlapping QTL from higher-level taxonomies. Results with adj. *p*-value < 0.05 (statistically significant) are bolded. Permutations were calculated only for peaks with the lowest likelihood *p*-value in regions with peak overlaps. OTU numbers are assigned by Greengenes database, ‘NR’ prefixes denote “New Reference” OTUs defined as those with sequences that failed to match the reference and are clustered *de novo*. ‘NCR’ prefixes denote “New Clean-up Reference” OTUs that failed to match the new reference OTU collection and are assigned a new random number. Complete table of QTL results for OTUs can be found in **Tables S6**.

| Taxa |  | chr<sup>a</sup> | maxlod<sup>b</sup> | pos<sup>c</sup> | from<sup>d</sup> | to<sup>e</sup> | *p*-value | adj.*p*-value | perm.*p*-value |
| --- | --- | --- | --- | --- | --- | --- | --- | --- | --- |
|  | OTU 421792 (order Bacteroidales) | 5 | 5.52 | 118.69 | 118.67 | 118.79 | 6.36E-04 | 0.088 | NA |
|  | OTU 460953 (order Bacteroidales) | 5 | 5.71 | 118.67 | 118.63 | 118.74 | 4.49E-04 | 0.085 | NA |
|  | OTU 190835 (order Bacteroidales) | 5 | 5.52 | 118.67 | 118.67 | 118.69 | 6.35E-04 | 0.090 | NA |
|  | OTU 209408 (order Bacteroidales) | 5 | 6.84 | 118.67 | 118.52 | 118.82 | 5.02E-05 | 0.066 | 0.195 |
|  | OTU NR.OTU100 (family Ruminococcaceae) | 5 | 8.35 | 31.93 | 30.25 | 32.10 | 2.46E-06 | 0.052 | 0.015 |
|  | **OTU 338796 (family Ruminococcaceae)** | **2** | **8.28** | **170.56** | **169.64** | **171.00** | **2.84E-06** | **0.032** | **0.019** |
|  | OTU NR.OTU95 (family Ruminococcaceae) | 5 | 7.65 | 31.83 | 31.30 | 32.00 | 1.01E-05 | 0.065 | NA |
|  |  | 5 | 7.45 | 32.27 | 32.11 | 32.39 | 1.51E-05 | 0.065 | NA |
|  | OTU 336810 (family Ruminococcaceae) | 2 | 6.47 | 170.54 | 170.27 | 170.59 | 1.03E-04 | 0.093 | NA |
|  | **OTU NCR.OTU170146 (family Ruminococcaceae)** | **5** | **7.68** | **33.33** | **32.27** | **35.85** | **9.58E-06** | **0.026** | **0.056** |

<sup>a</sup>Chromosome in which lies the QTL
<sup>b</sup>Maximum LOD score within the QTL
<sup>c</sup>Position in the chromosome (Mbp) where the maximum LOD score is found
<sup>d</sup>Chromosomal position (Mbp) where the QTL begins
<sup>e</sup>Chromosomal position (Mbp) where the QTL ends

For every bacterial taxon from the five taxonomic levels with a statistically significant QTL association, we mapped the OTUs belonging to that taxon. We applied a 50% zero cut-off filter to only retain common OTUs and generated QTL mappings and assessed significance as described above for the five taxonomic levels.

When necessary for comparison, genomic coordinate spans from other publications were translated from human hg19 assembly to mouse mm10 assembly using LiftOver (UCSC). Particularly in the case of small spans or single nucleotides, LiftOver might require expanding the window being mapped. In our case, we iteratively increased the window by adding a padding of 0, 10,100,1000,10000, and 100000 bps on each side of the region of interest until a mapping was achieved. All mapped entries listed include the final span of the genomic coordinates used including padding (**Table S7D**).

### Gene Set Pathway Analysis

We used Ingenuity Pathway Analysis (IPA®, QIAGEN Redwood City, CA) software to conduct gene set pathway analysis on the protein-coding non-predicted genes within our QTL regions. Genes were uploaded as NCBI Gene IDs for ease of mapping across IPA’s source databases. All analyses were constrained to consider only direct relationships and exclude any annotation predictions. Additionally, we used IPA’s stringent filter to constrain the analysis to Mouse annotation while considering all Tissues and Cell Lines. IPA by default shows uncorrected *p*-values for enrichment analyses. We customized all charts and tables to indicate Benjamini-Hochberg False Discovery Rate instead. In total, we submitted 6 gene lists for parallel analyses, all of which were filtered to exclude predicted genes and non-protein coding genes: (1) all genes within any significant QTL region at any taxonomic level, (2) Bacillales only, (3) Bacteroidales only, (4) Mollicutes only (5) Ruminococcaceae only, and (6) *Staphylococcus* only. Many taxonomic groups result in similar-enough QTLs that their gene sets are identical, these groups are the result of picking the lowest taxon for any identical gene sets while covering all the taxa studied (for instance, phylum Tenericutes is excluded as it matches the results of class Mollicutes).

### Data Availability

Our study was performed on a subset of Diversity Outbred mice from the Allan Pack Sleep Study; the genotypes can be downloaded from the Jackson Lab Diversity Outbred Database (DODB) website (https://dodb.jax.org). QIIME demultiplexed fastq files with microbiome data are available in the NCBI SRA, BioProject ID: PRJNA639769 (https://www.ncbi.nlm.nih.gov/bioproject/639769). All Supplemental Materials (**File S1**, **Figures S1-S5**, and **Tables S1-S9**) have been uploaded to GSA FigShare under “Supplemental Material for Schlamp et al., 2020”, and a description of each can be found at the end of the manuscript.

## RESULTS

### Variation of gut microbiota

High-throughput sequencing of fecal samples from 247 three month old male mice from the Diversity Outbred Mouse Panel generated 15,149,384 16S rRNA gene sequences that passed the quality filtering criteria after demultiplexing (see **Materials and Methods**). On average, 61,334 sequences were obtained per sample (ranging from 17,658 to 135,803 sequences). Sequences were sorted into 57,014 operational taxonomic units (OTUs) at 97% identity against the Greengenes 8_13 database using open-reference OTU picking (**Table S1A**). Next, OTUs were summarized at five levels of taxonomy (phylum, class, order, family, genus) (**Table S2A**). In order to focus on the most abundant microbes, only the taxa present in at least 50% of samples (i.e. present in 124 samples or more) were used for all following analysis, leaving a total of 75 taxa to test at the five levels of taxonomy (6 phyla, 8 classes, 11 orders, 20 families, and 30 genera). The most predominant taxa at the phylum level were Firmicutes (average relative abundance = 48.64%) and Bacteroidetes (46.41%), which is consistent with previous findings in mice (Benson *et al.* 2010; Mcknite *et al.* 2012; Org *et al.* 2015). The relative abundances of these taxa were highly variable, with Firmicutes ranging from 11% to 94%, and Bacteroidetes ranging from 1% to 88% (**Figure 2, Figure S1**).

**Figure 2.**
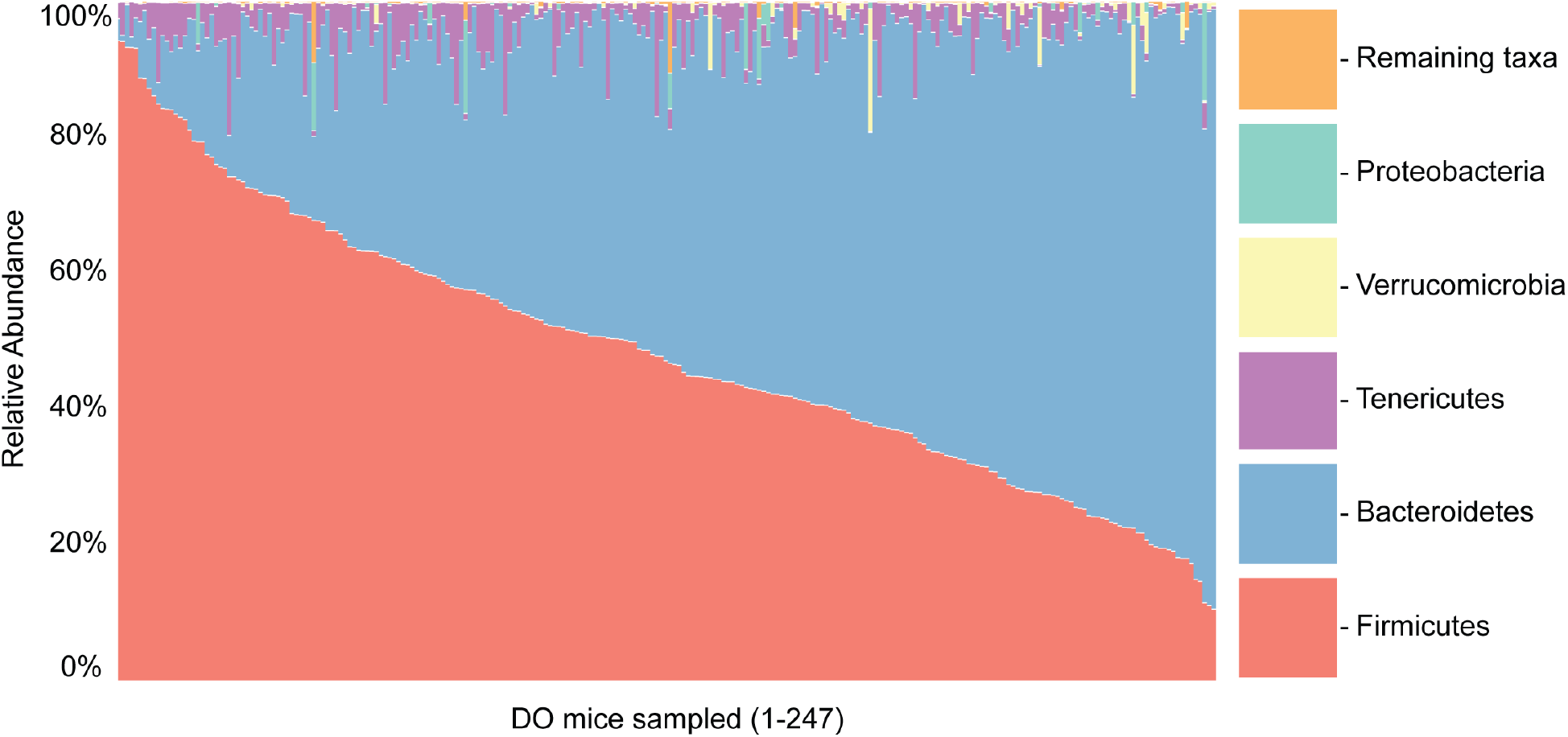
Relative abundances of top ten most abundant phyla across the 247 DO mice. Relative abundances shown, mouse samples sorted by phylum Firmicutes, the most abundant phylum.

The top 8 most abundant genera were present in at least 99% of the samples. The two most abundant genera were an unidentified genus within Bacteroidales family S24-7 (average relative abundance = 43.89%, ranging from 1% to 88%) and another unidentified genus within Clostridiales (32.35%, ranging from 4% to 78%), consistent with previous findings in mice (Shin *et al.* 2016).

When dealing with uneven sequence counts across samples, microbiome studies commonly normalize the data by rarefying sequence counts, which consists of randomly selecting from each sample an equal number of sequences without replacement (Weiss *et al.* 2017). It has been argued, however, that rarefaction is not an ideal approach due to valuable data being discarded (Mcmurdie AND Holmes 2014). Therefore, we decided to present our analysis of the non-rarefied data using sequence counts per sample as a covariate, noting also that the rarefied data consisted of highly similar relative abundances, and provided similar heritability and QTL results (see **File S1** for a detailed breakdown of these metrics).

### Heritability estimation

Each of the 247 individual DO mice used in this study represents a unique genomic combination of alleles from the original eight progenitor lines. The unit of inference for phenotypes was the rank Z-score transformed relative abundance of each taxon at each taxonomic level (phylum, class, order, family, genus) in each individual mouse, while the units of genetic inference were the SNP genotypes at each of 57,973 sites for each mouse using the MegaMUGA mouse genotyping array (**Figure 1**). Each SNP genotype is represented by an eight-founder state probability that corresponds to the probabilities contributed by each founder at each SNP (Svenson *et al.* 2012) and those eight-founder probabilities are used to fit the linear models (see **Material and Methods**).

We estimated narrow-sense “SNP” heritability (*h^2^*) using a linear mixed model in R-package *lme4qtl* (Ziyatdinov *et al.* 2018). A linear mixed model was used to predict whether the effects of the autosomal genotype on the phenotype is proportional to the genetic similarity between the mice, after adjustment for known factors. Thus, calculations were based on the kinship matrix (genetic similarity; also called genetic relatedness matrix (GRM)), expression of a phenotype (taxon relative abundance) across all samples, and additional covariates (such as sequencing run, read counts, and cage effect). Significance was assessed by a restricted likelihood ratio test using R-package *RLRsim* (Scheipl *et al.* 2008). More details can be found in **Materials and Methods**. Heritability estimates ranged from 0% to 40%. In total, 23 of the 75 tested taxa were significantly heritable (RLRT *p*-value < 0.05); we additionally note multiple-hypothesis normalized Benjamini-Hochberg (BH) False Discovery Rates (**Figure 3**, **Table S3A**). We hypothesized that higher-level taxa would be found to be more heritable than lower level taxa, assuming strong functional relatedness between members of the same taxonomic group, but found that there is no consistent trend between the taxonomy level and heritability. Our most heritable taxon, the class Mollicutes (40%, RLTR *p*-value of 0.0017, BH *p*-value of 0.0884) had a higher heritability estimate than any clade below it (genus *Anaeroplasma* with 29% and an unclassified genus in order RF39 with 35%). In contrast, the heritability estimate of class Bacilli (25%), is surpassed by its subclade, the order Lactobacillales (33%), which is in turn surpassed by its genus *Lactobacillus*, our second most heritable taxon (36%, RLTR *p*-value of 0.0076, BH *p*-value of 0.1035). Proportion variance estimates for kinship and cage for all taxa and their taxonomic level are presented in **Figure 3**.

**Figure 3.**
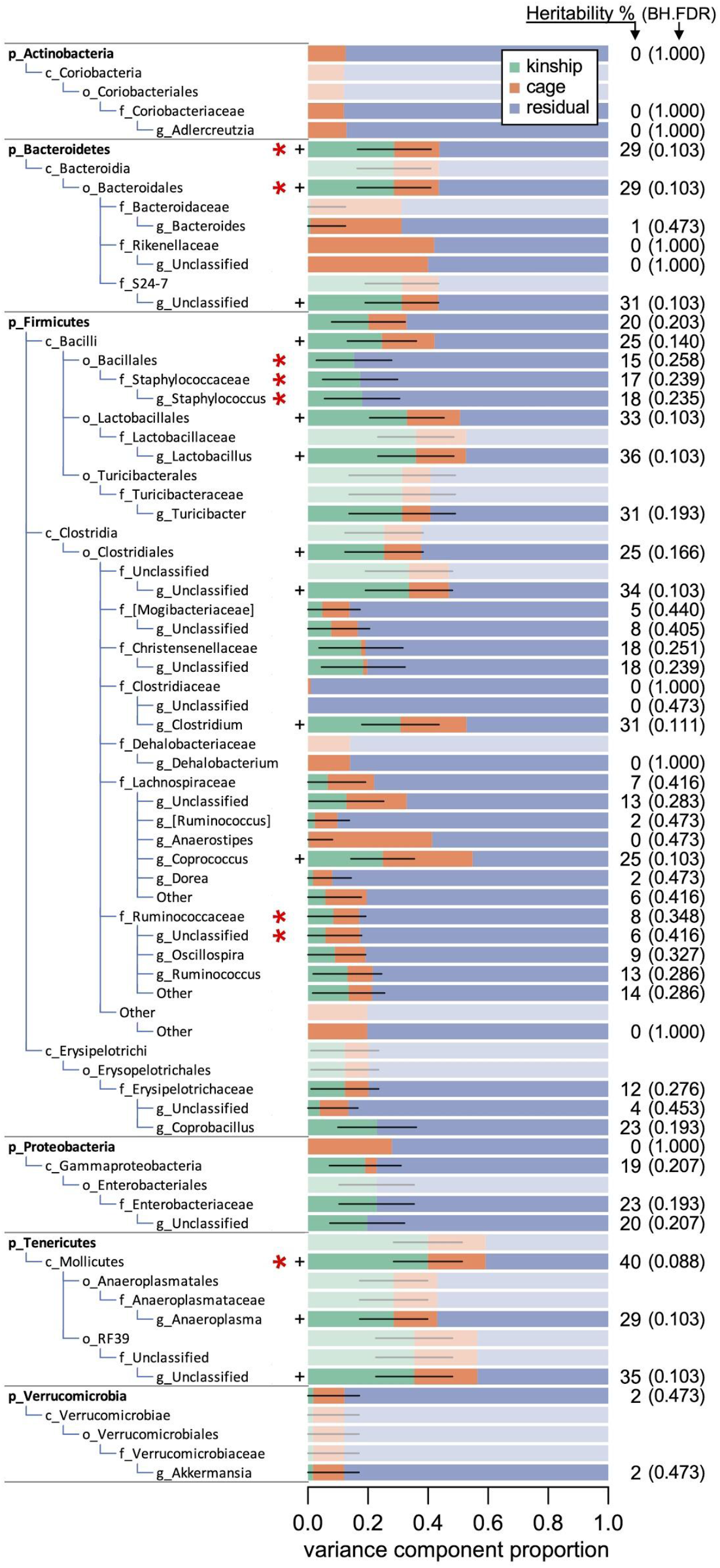
Proportion of variance estimates for kinship and cage for all taxa. Proportion of variance estimates for kinship (green), cage effects (orange), and unexplained residual effects (blue) for each taxon. The kinship proportion of variance is an estimate of narrow sense heritability. Heritability percentages are shown on the left. Heritability standard errors are shown with black horizontal lines. Designations p_, c_, o_, f_, and g_ are for phylum, class, order, family, and genus, respectively. When results are identical across taxa in the same phylogenetic branch, only the lowest (most specific) taxa are shown and the rest are shaded out. Heritability significance is marked with one plus (**+**, RLTR *p*-value < 0.05) and BH FDR is shown in parentheses next to heritability percentages. Taxa marked with a red asterisk have statistically suggestive QTL (⭑, adj. *p*-value < 0.1). Complete table of heritability results, including rarefied data, can be found in **Table S3**.

### QTL Mapping

QTL mapping of the bacterial taxa at the five taxonomic levels revealed findings that suggest statistically significant associations between host genotype and relative abundances of certain taxa. QTL regions on autosomes were found using the R-package *lme4qtl* (Ziyatdinov *et al.* 2018). Significance was assessed first by comparison of models with and without genotype via a likelihood ratio test, followed by a genome-wide permutation test. The reported *p*-values were corrected for multiple testing across SNPs (but not across taxa). In total, genetic associations with the abundance of family Ruminococcaceae, family Staphylococcaceae, and genus *Staphylococcus* were found to be statistically significant (adj. *p*-value < 0.05), and additional genetic associations with phylum Bacteroidetes, order Bacteroidales, order Bacillales, and class Mollicutes were statistically suggestive (adj. *p*-value < 0.1). QTLs of order Bacteroidales, genus *Staphylococcus*, and family Ruminococcaceae were also statistically suggestive at a permutation *p*-value < 0.1 (**Table S5**).

QTL regions are defined by all contiguous SNPs with LODs above significance threshold of adjusted *p*-value < 0.1, as illustrated in **Figure 4C**. Multiple QTL for various taxa overlapped with the QTL regions for their parent taxa, such as a QTL hit for genus *Staphylococcus* (a genus in the family Staphylococcaceae) overlapping the QTL hit for family Staphylococcaceae (**Table 1**). The relationship between loci and microbial abundance is treated as an independent association analysis per taxa. An example of how these parallel analyses detect similar genomic regions across related taxa is further illustrated in **Figure 5**.

**Table 1.** QTL regions for taxa at five taxonomic levels. Only showing ranked results with adj. *p*-value < 0.1 (statistically suggestive). Results with adj. *p*-value < 0.05 (statistically significant) are bolded. When results were overlapping across taxa in the same phylogenetic branch (such as phylum Bacteroidetes and order Bacteroidales), permutations were calculated only for the lowest (most specific) taxon. Complete table of QTL results, including rarefied data, can be found in **Table S4**.

| | | chr<sup>a</sup> | maxlod<sup>b</sup> | pos<sup>c</sup> | from<sup>d</sup> | to<sup>e</sup> | $p$-value | adj.$p$-value | perm.$p$-value |
| --- | --- | --- | --- | --- | --- | --- | --- | --- | --- |
| Taxa | phylum Bacteroidetes | 5 | 7.65 | 118.58 | 117.39 | 118.82 | 1.02E-05 | 0.068 | NA |
|  | order Bacteroidales | 5 | 7.67 | 118.58 | 117.39 | 118.82 | 9.69E-06 | 0.065 | 0.042 |
|  | order Bacillales | 19 | 8.24 | 27.02 | 26.55 | 27.42 | 3.12E-06 | 0.055 | NA |
|  |  | 19 | 6.95 | 27.82 | 27.82 | 27.97 | 4.02E-05 | 0.085 | NA |
|  | family Staphylococcaceae | **19** | **8.09** | **27.04** | **26.55** | **27.46** | **4.17E-06** | **0.032** | NA |
|  |  | **19** | **7.84** | **27.82** | **27.61** | **28.18** | **6.96E-06** | **0.032** | NA |
|  |  | 19 | 6.58 | 32.10 | 31.83 | 32.22 | 8.33E-05 | 0.092 | NA |
|  |  | 19 | 6.45 | 32.43 | 32.43 | 32.43 | 1.07E-04 | 0.098 | NA |
|  | genus *Staphylococcus* | **19** | **8.26** | **27.04** | **26.51** | **27.46** | **3.00E-06** | **0.025** | **0.019** |
|  |  | **19** | **7.93** | **27.82** | **27.61** | **28.20** | **5.78E-06** | **0.025** | **0.033** |
|  |  | 19 | 6.64 | 32.10 | 31.83 | 32.22 | 7.39E-05 | 0.082 | 0.271 |
|  |  | 19 | 6.53 | 32.43 | 32.43 | 32.46 | 9.14E-05 | 0.082 | 0.304 |
|  | family Ruminococcaceae | **2** | **7.01** | **170.57** | **170.51** | **170.66** | **3.63E-05** | **0.039** | **0.137** |
|  |  | 2 | 6.30 | 170.71 | 170.71 | 170.71 | 1.44E-04 | 0.100 | 0.427 |
|  |  | **5** | **7.18** | **31.93** | **31.77** | **32.19** | **2.59E-05** | **0.035** | **0.106** |
|  |  | **5** | **7.44** | **32.52** | **32.27** | **33.36** | **1.55E-05** | **0.035** | **0.066** |
|  | Unclassified genus in family Ruminococcaceae | 2 | 7.89 | 170.56 | 170.51 | 170.64 | 6.22E-06 | 0.090 | NA |
|  | class Mollicutes | 1 | 7.00 | 121.32 | 120.23 | 125.20 | 3.66E-05 | 0.089 | 0.142 |

<sup>a</sup> Chromosome in which lies the QTL
<sup>b</sup> Maximum LOD score within the QTL
<sup>c</sup> Position in the chromosome (Mbp) where the maximum LOD score is found
<sup>d</sup> Chromosomal position (Mbp) where the QTL begins
<sup>e</sup> Chromosomal position (Mbp) where the QTL ends

**Figure 4.**
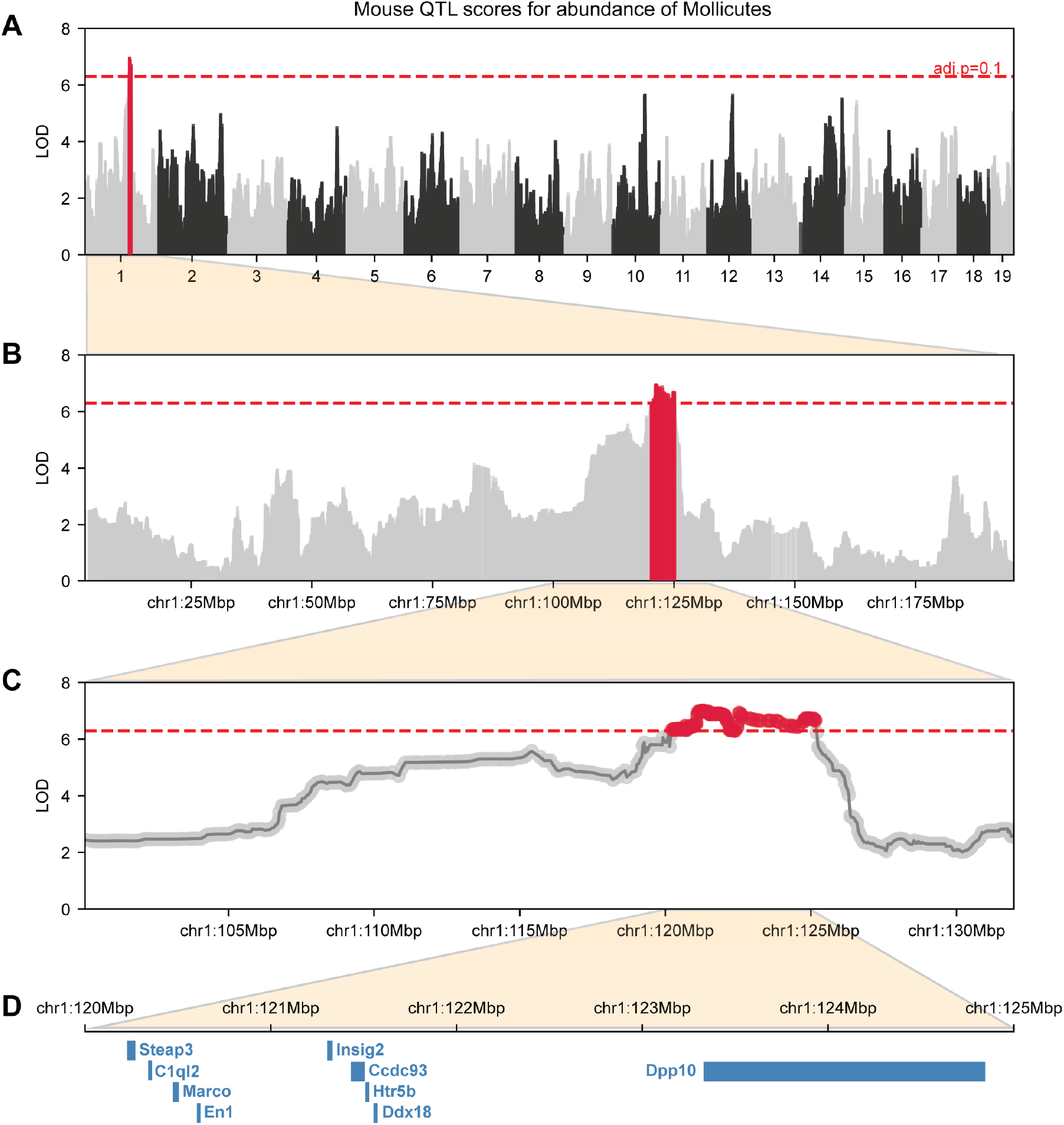
QTL scores for abundance of class Mollicutes. **(A)** LOD score profile of genome-wide QTL mapping for relative abundance of class Mollicutes shows a significant QTL region (in red) on chr1. Horizontal axis shows genome physical location by chromosome, vertical axis shows LOD score at each site. Horizontal dashed red line marks the significance threshold at adjusted *p*-value < 0.1. (**B**) Chr1 zoomed in shows the significant QTL region in red (chr1:120.24-125.15 Mbp). (**C**) Further zoom into the area of interest shows in clearer detail how the QTL region is determined by a collection of contiguous significant SNPs (above significance threshold). (**D**) *Mus musculus* protein-coding genes within the QTL region are colored in blue.

**Figure 5.**
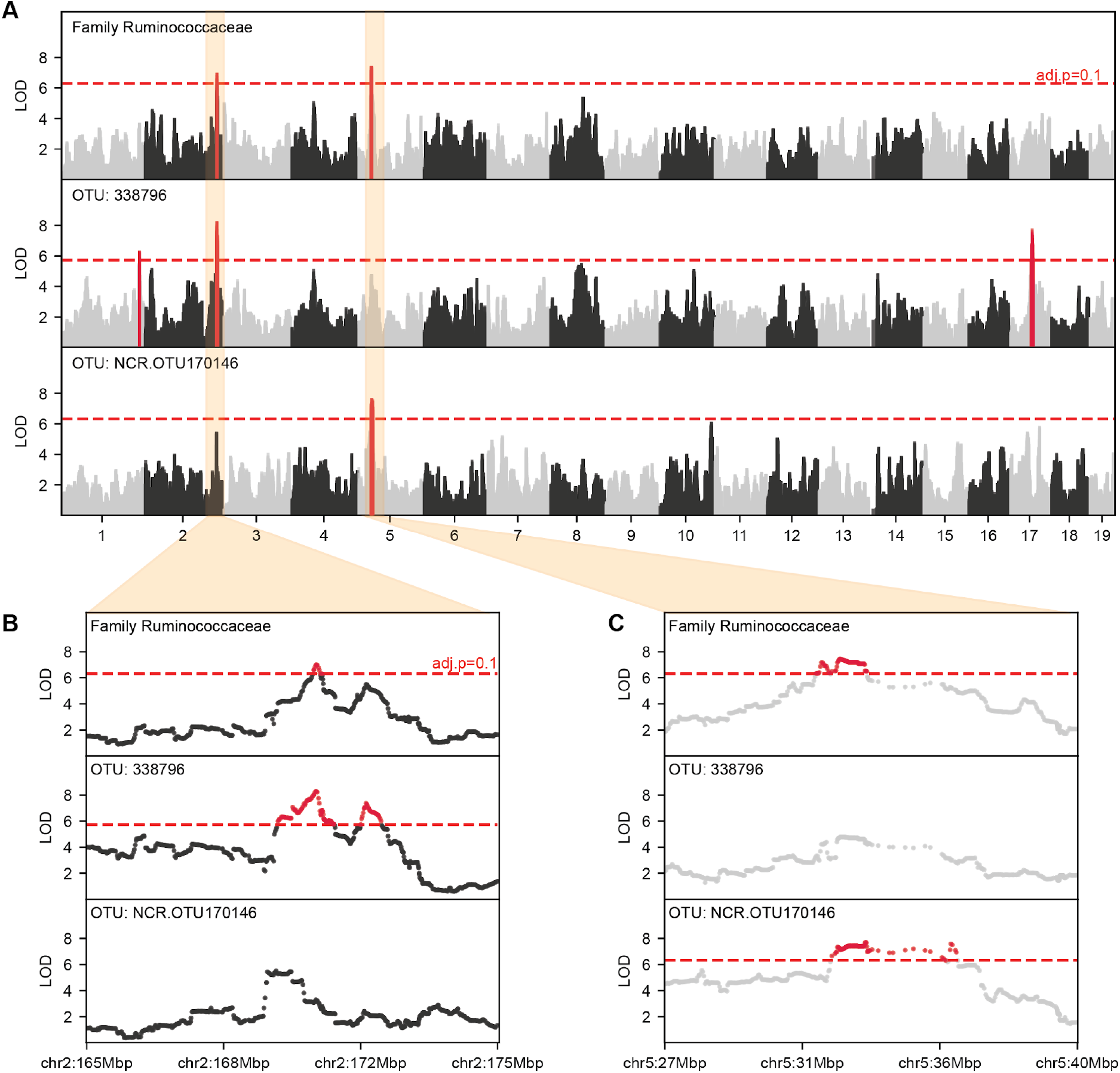
Overlap of QTL regions across taxa in the same phylogenetic branch. (**A**) LOD score profile of genome-wide QTL mapping for relative abundance family Ruminococcaceae (top panel), and two OTUs found within family Ruminococcaceae: OTU 338796 (middle panel), and NCR OTU 170146 (bottom panel). (**B**) Zoom into the area of interest in chr2 shows overlap in QTL regions between family Ruminococcaceae (170.51-170.66Mbp, top panel) and OTU 338796 (169.64-171.00Mbp, middle panel). (**C**) Zoom into the area of interest in chr5 shows overlap in QTL regions between family Ruminococcaceae (32.27-33.36Mbp, top panel) and NCR OTU 170146 (32.27-35.85Mbp, bottom panel). Horizontal axis shows genome physical location by chromosome, vertical axis shows LOD score at each site. Horizontal dashed red line marks the significance threshold at adjusted *p*-value < 0.1.

### OTU level analysis

Next, we decided to increase the specificity of the taxonomic classifications to operational taxonomic units (OTUs) by compiling all OTUs identified within taxa that had statistically suggestive QTL (**Table 1**). We filtered out OTUs that were present in less than 50% of the samples, resulting in 362 OTUs. QTL mapping performed on these selected OTUs resulted in 28 OTUs with at least one statistically significant association (adj. *p*-value < 0.05), and 33 additional OTUs with at least one statistically suggestive association (adj. *p*-value < 0.1) (**Table S6**).

QTL associations to OTUs sometimes overlapped with QTL regions associated to taxa at higher taxonomic levels, with the most significant ones corresponding to wider QTL regions (**Table 2**). These results are interesting because if the overlapping QTL region associated with the broader taxonomic group is narrower and more specific than the region seen on an individual OTU, this might suggest a cumulative effect of multiple sub-taxonomies driving a stronger signal at the broader taxonomic level. For example, QTL for OTU 338796 (chr2:169.64-171.00Mbp) and NCR OTU 170146 (chr5:32.27-35.85Mbp) within family Ruminococcaceae were both statistically significant (**Table 2**) and overlapped with QTL regions for Ruminococcaceae (chr2:170.51-170.66Mbp and chr5:32.27-33.36Mbp, respectively) (**Table 1**), but the QTL regions for the OTUs were both wider, as shown in **Figure 5**. Note that factors such as the local recombination intensity profile and SNP density will affect the width of the QTL regions equally across taxonomies, since the relative abundance of each taxon at each taxonomic level is considered as a single phenotype queried against the same static, underlying set of SNPs.

### Comparison to other studies

Results from other published studies on heritabilities of the various bacterial taxa in the gut microbiome of mice, pigs, and humans were compiled and compared with our results (**Figure 6, Table S7**). We find new evidence of heritability of bacterial taxa in mice only previously seen in human studies. For example, we observed significant heritability in the phylum Tenericutes as well as several of its subclades, including genus *Anaeroplasma* and order RF39. These results were consistent across both our rarefied and non-rarefied datasets, and had not been seen in any other mouse studies, either because they did not detect these taxa in their studies or their results failed to identify significant heritability. This novel result is similar to previous host-microbe associations seen in human studies where significant heritabilities for this taxonomic lineage were identified in phylum Tenericutes (*h^2^* = 0.34 (Goodrich *et al.* 2016) and 0.23 (Lim *et al.* 2017)), class Mollicutes (*h^2^* = 0.32 (Goodrich *et al.* 2016) and 0.23 (Lim *et al.* 2017)), and order *RF39* (*h^2^* = 0.31 (Goodrich *et al.* 2016)).

**Figure 6.**
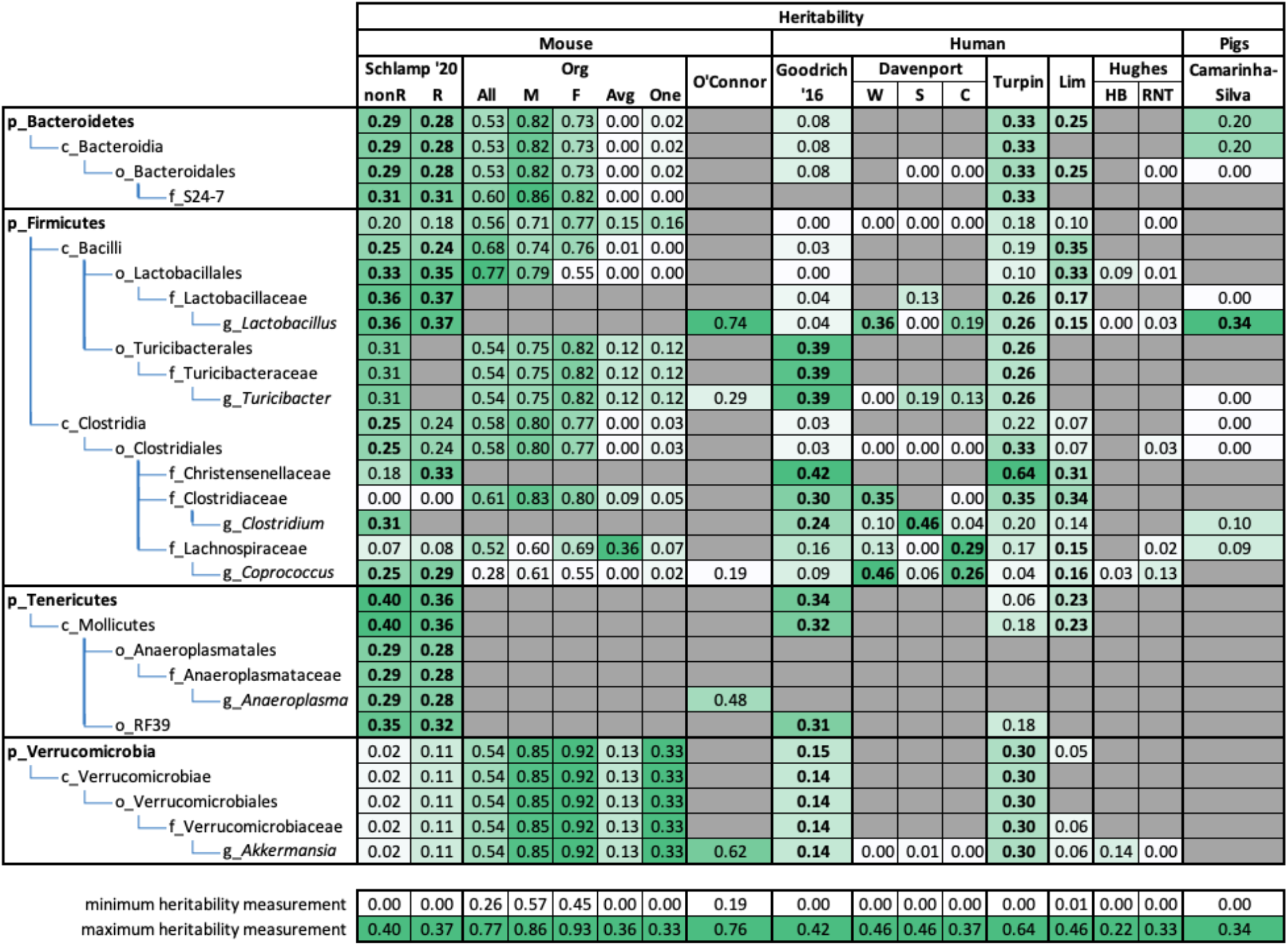
Comparison of taxon heritabilities across mouse, human, and pig studies. The green shading over heritability estimates ranges from each study’s lowest heritability estimate (white) to each study’s highest heritability estimate (green) to highlight the relative heritability of each taxa per study. Statistically significant results are shown in bold font when significance is reported. For our Diversity Outbred study, we report both non-rarefied (nonR) and rarefied (R) results. For Org *et al.* (2015) we report results using all mice (All), just males (M), just females (F), an average per strain (Avg), and a single mouse per strain (One). Org *et al.* (2015) and O’Connor *et al.* (2014) did not report significances. For Goodrich *et al.* (2016) the estimates are calculated by the ACE model, bold values indicate estimates with a 95% confidence interval not overlapping 0. For Davenport *et al.* (2015) the estimates are the proportion of variance explained (PVE) estimates (“chip heritability”), we report winter (W), summer (S), and combined seasons (C) datasets, and bold values indicate estimates with a standard error not overlapping 0. For Turpin *et al.* (2016) and Lim *et al.* (2017) estimates are polygenic heritability (H2r). For Camarinha-Silva *et al.* (2017) and Hughes *et al.* (2020) estimates are narrow-sense heritability (*h^2^*). Grey indicates that the taxon was not observed or excluded in a given study. Figure adapted from Goodrich *et al.* (2016). Comparisons relevant to the text are shown here, with the full comparison found in **Table S7**.

In some instances, taxa that we did not identify as being significantly heritable — and in fact have some of our lowest heritability scores — are reported to have high heritability in other studies. We show some examples of this in **Figure 6**: families Clostridiaceae and Lachnospiraceae as well as the entire phylum Verrucomicrobia. Interestingly, both of these families have significantly heritable subclades, whereas the entire branch of phylum Verrucomicrobia had low heritability estimates. We see a very low heritability estimate in both our non-rarefied and rarefied datasets for the genus *Akkermansia* (*h^2^* = 0.02) and every taxonomic level up to phylum Verrucomicrobia, yet estimates for mice in other studies were as high as *h^2^* = 0.92 (Org *et al.* 2015), and *h^2^* = 0.62 (O’Connor *et al.* 2014). This discrepancy between heritability estimates for *Akkermansia* is not mouse specific, as human microbiome studies see similarly conflicting results in their heritability estimates for this same genus: Reporting significantly high (*h^2^* = 0.30 (Turpin *et al.* 2016)), significantly low (*h^2^* = 0.14 (Goodrich *et al.* 2016)), and close to zero and not significant estimates (*h^2^* = 0, 0.01 (Davenport *et al.* 2015), and 0.06 (Lim *et al.* 2017)).

In addition to comparing our heritability estimates with other studies, we also contrasted our QTL mapping results of the gut microbiome with those from previous QTL and GWA studies (**Figure 7, Table S7**).

**Figure 7.**
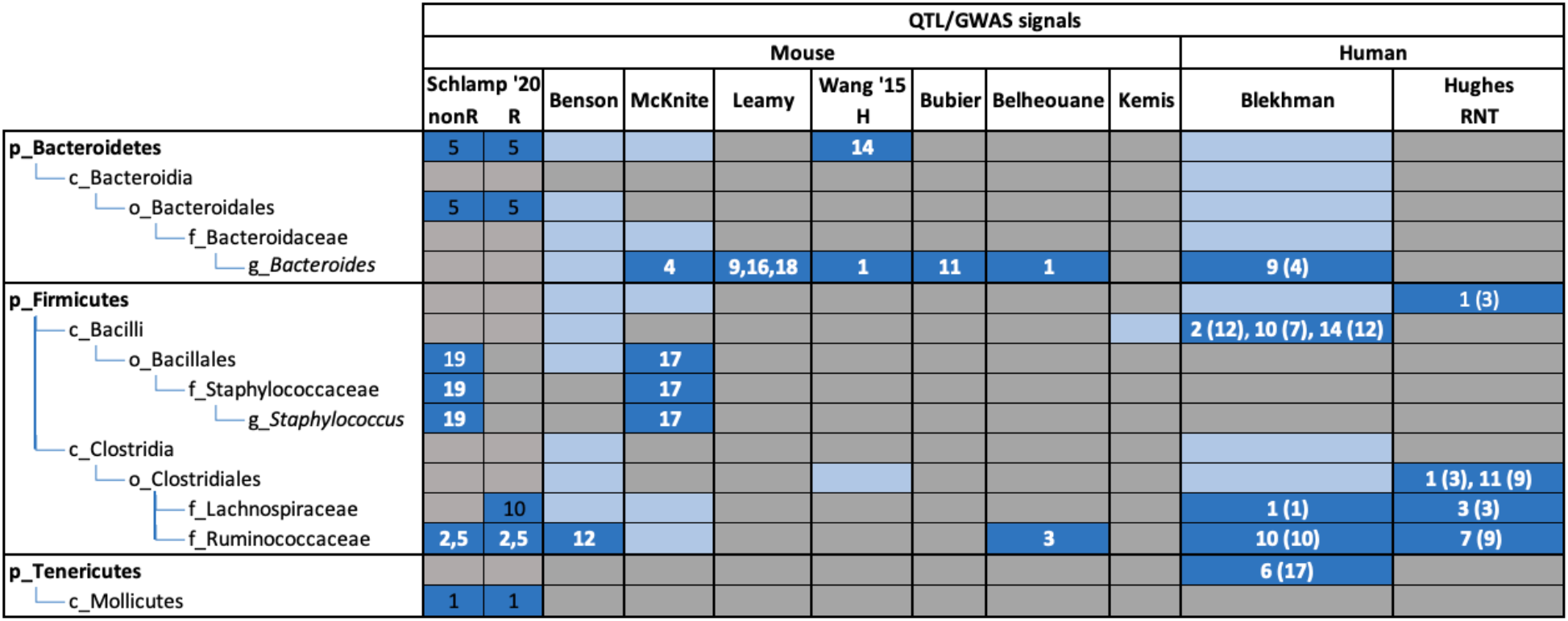
Comparison of taxa with QTL associations across mouse and human studies. Associations with each taxon are marked in dark blue if statistically suggestive and bolded in white if statistically significant, or light blue if not significant. Gray indicates that the taxon was not observed or excluded in a given study. The chromosome numbers where the QTL were found are denoted in each box. For our Diversity Outbred study, we report both non-rarefied (nonR) and rarefied (R) results. In the human studies, the corresponding syntenic mouse chromosome was added in parenthesis. Figure adapted from Goodrich *et al.* (2016). Selected comparisons shown, full comparison found in **Table S7**.

We identified statistically significant QTL associations for the order Bacillales as well as for the family Staphylococcaceae and the genus *Staphylococcus* within Bacillales in chr19; another mouse study also found statistically significant QTL associations for all of the same taxa but on chr17 (Mcknite *et al.* 2012). A human microbiome study found statistically significant QTL regions for the class Bacilli, which comprise the above mentioned order and families (Blekhman *et al.* 2015).

Family Ruminococcaceae has been previously found to have significant QTL associations both in mice (chr12 (Benson *et al.* 2010) and 3 (Belheouane *et al.* 2017)) and humans (Blekhman *et al.* 2015, Hughes *et al.* 2020). In our study, Ruminococcaceae was identified as associated with chromosomes 2 and 5. We also identified a QTL hit for the phylum Bacteroidetes in chr5 while another mouse study identified a significant hit in chr14 (Wang *et al.* 2015). Within Bacteroidetes, even though we did not find any significant QTL results for the genus *Bacteroides*, many other mouse studies have (chr1 (Wang *et al.* 2015, Belheouane *et al.* 2017), 4 (Mcknite *et al.* 2012), 9 (Leamy *et al.* 2014), 11 (Bubier *et al.* 2018), 16 (Leamy *et al.* 2014), and 18 (Leamy *et al.* 2014)) as well as a human study (Blekhman *et al.* 2015).

Phylum Tenericutes had a significant hit in chr1 in both our non-rarefied and rarefied datasets, and family Lachnospiraceae had a statistically suggestive QTL in chr10 in our rarefied dataset but not in our non-rarefied dataset. Both of these taxa had significant QTL hits in a human study (Blekhman *et al.* 2015).

Finally, we did not observe any QTL overlaps with Kemis *et al.* (2019) (**Table S7**), which also used the Diversity Outbred mice population in their microbiome association study. This lack of overlap is likely due to the highly different diet used in their experiments (high-fat, high-sucrose), and further highlights the strong impact of diet alone in microbiome composition.

### Gene level analysis

Examining the QTL mapping results from previous studies, it was apparent that although different studies might all have found significant QTL regions for a particular bacterial taxon, they identified different genomic positions as showing associations. In order to identify common functions and diseases associated with the genes within our QTL regions, we used Ingenuity Pathway Analysis (IPA®, QIAGEN Redwood City, CA) to run a cumulative gene set enrichment analysis on all 1423 genes associated with non-rarefied microbiome abundance spanning 7 significant QTL and 11 suggestive QTL across the five taxonomic levels (phylum, class, order, family, genus) (**Table S4**) and 54 significant QTL and 232 suggestive QTL at the OTU level (**Table S6**). All genes found within each QTL region were included. When QTL regions overlapped across taxa of the same phylogenetic branch (as illustrated in **Figure 5**), overlapping genes were only counted once. Additionally, we ran taxon-specific enrichment analysis to profile the specific functions and diseases associated with genes in the QTL regions associated with relative abundance of phylum Firmicutes (n = 23 genes), class Mollicutes (n = 9), order Bacteroidales (n = 10), family Ruminococcaceae (n = 15), and genus *Staphylococcus* (n = 8), excluding OTU-specific QTL regions.

Through the cumulative gene set analysis, we found 25 networks each containing subsets of our genes. We can try to characterize the biological significance of these networks by measuring the enrichment of disease and functional annotations in the genes of each network (**Table S8**). We find remarkable functional signatures in the highest-ranked of these 25 networks, with enrichment in the broad categories of Immunological Disease and Inflammatory Response (Network 1, **Figure 8A**), Lipid Metabolism and Molecular Transport (Network 2, **Figure 8B**), and Connective Tissue Development and Function (Network 3, **Figure 8C**). These associations are highly concordant with increasingly well-understood roles in host-microbiome interaction studies. In Network 1, we find the most enriched specific functions relate to microbiome associated phenotypes, namely hypersensitive reactions (BH-FDR = 7.56e-4), allergies (BH-FDR = 1.39e-3), and atopic dermatitis (BH-FDR = 6.35e-3). Additionally, we find that despite lack of overlap between the gene membership of these four highest-ranked networks, they all have a significant enrichment for functions in both Gastrointestinal Disease (BH-FDRs between 1.26e-3 and 2.47e-3) and Digestive System Development and Function (BH-FDRs between 3.41e-3 and 3.34e-2). Interestingly, we also find consistent significant enrichment of cancer annotations across all four networks with varying overlap of tissues: prostate and renal cancers (Network 1), metastasis and colorectal cancer (Network 2), and breast, ovarian, and gastrointestinal cancer (Network 3). Only liver cancer appeared to be enriched in all three networks. A full exhaustive list of significantly enriched categories, diseases, and functions can be found in **Table S9**.

**Figure 8.**
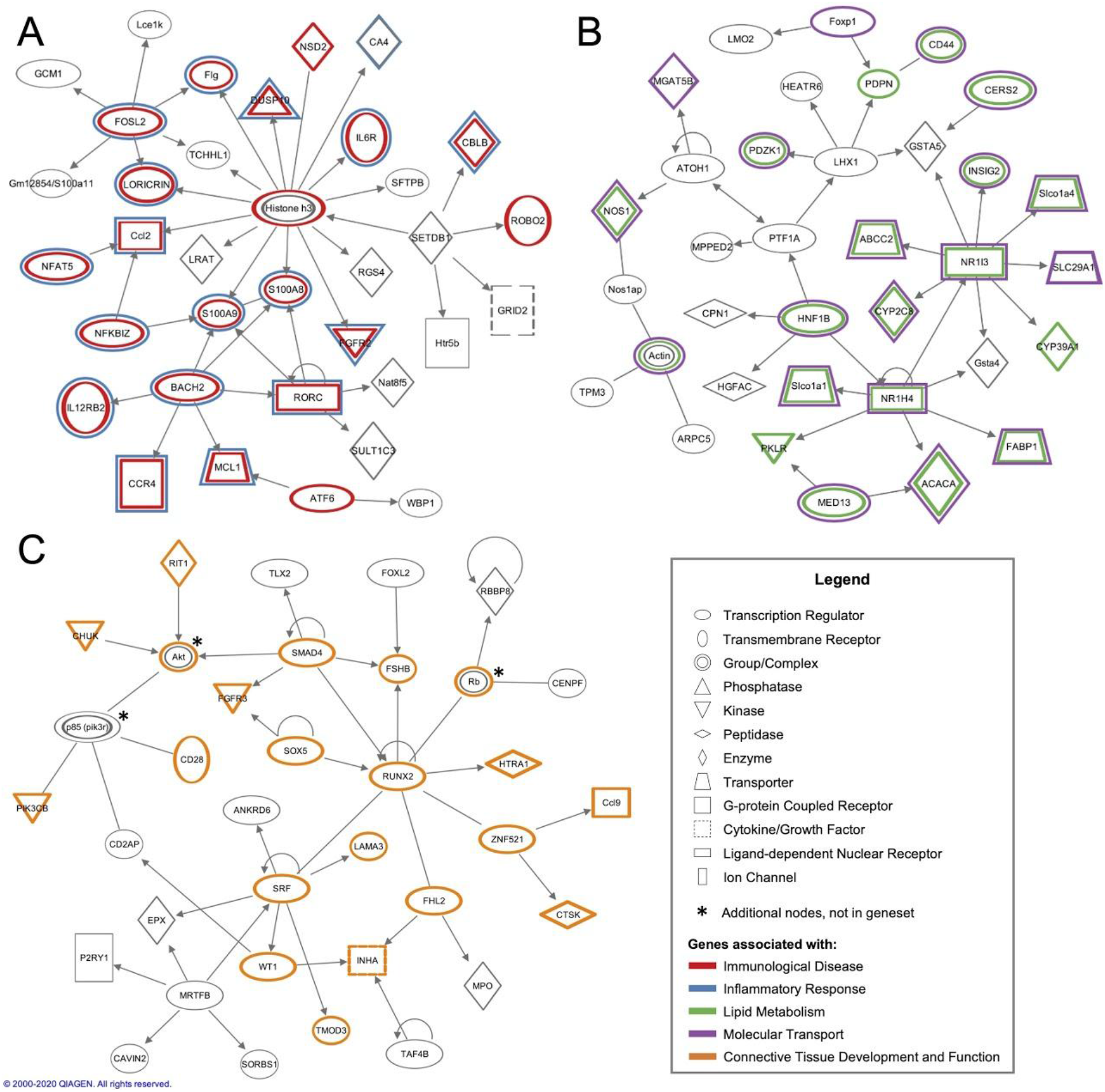
Ingenuity Pathway Analysis (IPA) three highest-ranked interaction networks generated from cumulative gene set analysis. Genes circled in color are all associated with disease and functional annotations as specified below. Nodes marked with an asterix belong to closely associated genes added by IPA that were not in the input dataset. (**A**) Network 1 shows genes associated with Immunological Disease (circled in red) and Inflammatory Response (blue). (**B**) Network 2 shows genes associated with Lipid Metabolism (green) and Molecular Transport (purple). (**C**) Network 3 shows genes associated with Connective Tissue Development and Function (orange).

Through taxon-specific enrichment analysis we find a consistent enrichment in development of adenocarcinoma (FDRs between 0.00% and 2.87%) in what are otherwise heterogeneous functional profiles (**Figure S2A-E**). We observe an enrichment of lipid metabolism pathway annotation through genes ASAH2, VLDLR, and SGMS1 in the phylum Firmicutes (FDRs 0.31% to 2.73%) all of which are again detected in its subclade, the genus *Staphylococcus* (FDRs 0.22% to 2.98%) (**Figure S2A,B**). We also observe an enrichment in breast and ovarian cancer annotations in phylum Firmicutes (FDRs 0.31% to 2.73%), which is shared with its larger subclade, the family Ruminococcaceae (FDRs 0.34% to 2.95%) (**Figure S2A,C**).

Finally, in both the cumulative and taxon-specific gene sets we find genes canonically tied to the commensal microbiome and pathogen-host interactions. The gene MARCO, which lies within a QTL for the abundance of class Mollicutes, encodes a pattern recognition receptor which is part of the innate antimicrobial immune system, binding both Gram-positive and Gram-negative bacteria. We consistently find genes associated with bacterial response in these gene sets, including TLR2, a membrane protein that recognizes bacterial, fungal and viral molecules and has been shown to have benign associations when binding a protein produced by the gut microbiome (Ottman *et al.* 2017). These gene sets also include both FCGR1A and FCER1G, fragments of the high affinity IgE Receptor, Spondin2, a cell adhesion protein that binds directly to bacteria and their components as an opsonin for the macrophage phagocytosis of bacteria, BPIFA1, an antimicrobial protein that inhibits the formation of biofilm by Gram negative bacteria, and both PGLYRP3 and PGLYRP4, both peptidoglycan recognition proteins that bind to murein peptidoglycan of Gram-positive bacteria.

## DISCUSSION

There exists a complex and multifaceted relationship between the gut microbiome and its host’s genome, where recent studies are beginning to show the true magnitude of these connections. Our results seek to further understand this relationship by measuring the heritability of bacterial relative abundance phenotypes and by categorizing the functional and disease pathways that may be associated with specific bacterial abundances in the mouse gut microbiome.

We detect the first instance of statistically significant heritability in the phylum Tenericutes in a mouse model. Specifically, we see that its subclade, class Mollicutes, is our most heritable taxon (40% BH *p*-value of 0.088). Dramatic increases in Mollicute abundance have been observed in mice when subjected to a high-fat, high-sugar diet in comparison to a plant polysaccharide-rich diet. This Mollicute bloom seems to come at the expense of Bacteroidetes abundance and an overall lower diversity in murine microbiomes (Turnbaugh *et al.* 2017). Understanding the heritable aspects of Mollicute abundance could help elucidate the host-genetic determinants of body weight control and the etiology of obesity, which has been thus far extremely challenging with host-genome GWAS alone (MÜLLER *et al.* 2018, Speakman et al. 2018).

Our second most heritable taxon is genus *Lactobacillus* (36% BH *p*-value of 0.103), which shares a similarly strong but benign association to body weight control in the literature. The genus *Lactobacillus* contains several species with strains commonly used as probiotics. In contrast to the Mollicute lineage, *Lactobacilli* have been used in mouse models of hyperlipidemia to show an increase of abundance of Bacteroidetes and Verrucomicrobia, and improving their lipid metabolism (Chen *et al.* 2014). The function of these clades as a whole is, however, not clear-cut: *Lactobacillus* is a large genus containing species and strains with differing roles and probiotic effects in humans (Mcfarland *et al.* 2018) and members of the class Mollicutes may have strain-specific positive effects on gastrointestinal disease in mice, rather than a negative phenotype as a whole (Zhai *et al.* 2019).

These examples present a microbiome-host interaction landscape in which associations between host health and microbiome abundance can be extremely taxon-specific, displaying functional heterogeneity at the species level. Building a baseline understanding of the resolution at which genetic associations change for different lineages is vital to build an understanding of health, function, and coevolution in microbe-host models.

We perform parallel analyses to find specific associations between genetic loci and individual taxonomic groups, treating sub-clades as independent phenotypes from their parent taxa during QTL calculations. This setup allows us to contextualize significant QTL across the bacterial taxonomy, and we find similarities in both the genomic regions detected and the functional annotation of covered genes for taxa in the same clade. We also greatly benefit from the type of QTL analysis facilitated by the DO mouse model, where the genotype for specific loci can be calculated and contrasted consistently across all samples, eliminating the need for the windowed confidence intervals which are common in this type of analysis.

We find functional associations in the gene sets identified by the QTL results that span disease and development phenotypes beyond obesity. We find QTL regions spanning genes with annotations for various phenotypes that are already widely studied in the context of host-microbiome interactions. Among them we see cancer-associated annotations, both in the more obvious gastrointestinal categories like colorectal cancer (Chen *et al.* 2012; Ahn *et al.* 2013, Zackular *et al.* 2014; Ericsson *et al.* 2015) and the surprising, but well-studied, breast (Yang *et al.* 2017; Fernández *et al.* 2018; Zhu *et al.* 2018), ovarian (Xu *et al.* 2020), and liver cancers (Yu AND Schwabe 2017). We also see an expected plethora of immune and inflammatory pathways, including some microbiome-associated disease hallmarks, including colitis (Knox *et al.* 2019), allergic response (Pascal *et al.* 2018), and atopic dermatitis (Kim AND Kim 2019). Beyond pathology, we see an enrichment of lipid metabolism pathways, coherent with the gut microbiome’s direct and indirect role in host lipid modulation (Ghazalpour *et al.* 2016; Heaver *et al.* 2018; Brown *et al.* 2019; Johnson *et al.* 2019).

The relationship between a host’s health and their microbiome seems increasingly complex. Links to host development, disease, and metabolism are still being found across body sites and a wealth of bioinformatic and modelling strategies continue to emerge (Malla *et al.* 2019). These results are a promising and heavily funded target for precision medicine (Proctor *et al.* 2019), identifying potential biomarkers for predisposition to type 1 diabetes (Uusitalo *et al.* 2016) and asthma (Durack *et al.* 2018) in children, and colorectal cancer in adults (Shah *et al.* 2018). As we move forward to understand the mechanisms underlying host-health modulation by the microbiome, it is imperative that we understand which parts of host genomes might have underlying associations with microbial species, both to understand the limitations of animal models as a relevant human proxy, and to determine whether host genetics plays a causal role.

Currently, there is a scarcity of studies discussing heritabilities and QTL mappings of bacteria within the gut microbiome. Despite the potential and funding of this field, there is still an absence of a standardized methodology for performing these studies that leads to the use of different procedures and analytical methods, making it increasingly difficult to compare results across studies (Goodrich *et al.* 2017; Kurilshikov *et al.* 2020). We see this in our comparisons of results with previous studies, as we do not observe consistent overlap in the estimated heritabilities and QTL associations in any one taxa. Depending on the study, we see differences in which covariates are able to be included, which databases or mapping algorithms are used to determine OTUs, and the manner in which results are reported. One salient example is our use of both the kinship matrix and co-housing as a random effect in our analysis, which required a tailored approach that extended the standard DO mice pipeline, which usually only allows a kinship matrix as random effect. Ultimately, the current state of the field for profiling different characteristics of the gut microbiome is still rapidly evolving and as it matures and more studies are undertaken, it will become easier to compare, validate, and aggregate results.

Although our results support the claim that host genetics can impact the gut microbiome composition in ways that are relevant to the health of the host, our study has some limitations. There is significant room for improvement in the statistical power of this study design through an increase in sample size (currently *n* = 247 DO mice).

Conducting QTL mapping with small sample sizes may lead to the ‘Beavis effect’ which is a failure to detect QTL of small effect sizes as well as an overestimation of effect size of the QTL that are discovered (Miles AND Wayne 2008). Our study is also subject to the trade-offs inherent in the Diversity Outbred design: since the genome of each mouse is a unique mixture of the 8 strains from the CC population, the genotype of each DO mouse is independent from other DO mice, and is irreproducible. This hampers the ability to generate biological replicates relative to inbred models, which in turn makes replicating results from the DO population limited to replication of marginal genetic effects. However, this limitation can be partially circumvented by using the CC lines as a form of validation, since they can provide reproducible genotypes (Svenson *et al.* 2012). Finally, associations between host genetics, microbiome abundance, and functional pathways must be investigated experimentally to confirm mechanism and causality. This is particularly difficult in the overlap of microbiome and genetic association, as hypothesis generation is a challenging and often gene-specific approach which must account for variation in both host and microbial communities.

Our results provide insight into the complex interplay between host genetics and the gut microbiome, and isolate associations between microbial taxa and QTL. Overall, this is a challenging analytical setup as we are trying to associate locus-specific variation with several inter-dependent phenotypes in a system with several covariates. Microbiome analyses are very sensitive to the traits, population, and environment under study, which we mitigate by taking into account co-housing and relatedness while also performing computationally intensive permutation tests to provide empirical *p*-values on our most significant QTL hits. As it stands, this method could be further utilized in a study with a novel microbial colonization (or other microbiome perturbation), where measuring the same phenotypes, in a similar setup, could be used to estimate the heritability and identify QTL for the successful introduction of a new taxon (or response to some other perturbation).

While most of the variation in the gut microbiome composition is not due to genetics but rather environmental factors (Rothschild *et al.* 2018), attributes of the gut microbiome that are clearly heritable may provide important insights about host-microbiome interactions and the mechanisms that impact microbiome composition. As the microbiome field moves toward novel disease models, biomarkers, and treatments, it is imperative that we understand the host-genetic variation that might influence the appropriateness of our models, the accuracy of our biomarkers, and the efficacy of new treatments.

## ACKNOWLEDGMENTS

The authors want to thank Noah Clark, Jessica L. Sutter, Qiaojuan Shi, Emily Davenport, and Afrah Shafquat for all the help and advice provided. F.S. was supported by a Presidential Life Science Fellowship (PLSF) from Cornell University. This work was supported by NIH grant R01 GM 070683.

## AUTHOR CONTRIBUTIONS

F.S., A.G.C, G.A.C, and R.E.L. conceived the study. A.P. provided the samples. F.S. extracted and generated the 16S rRNA gene sequencing data. F.S., E.J.C., P.S., J.K.G., R.E.L., G.A.C, and A.G.C. conceived the computational and statistical analyses. F.S., D.Y.Z., E.J.C., J.F.B., M.E., and P.S. performed the computational and statistical analyses. F.S., J.F.B., D.Y.Z., E.J.C, and A.G.C. wrote the manuscript.

## SUPPLEMENTAL MATERIAL

**File S1 - Analysis on rarefied data.** Detailed breakdown of variation of gut microbiota, heritability estimates, and QTL association results for rarefied data.

**Figure S1**- **Taxa relative abundance frequencies.** Stacked bar plots and box plots depicting relative abundance frequencies of the top ten most abundant taxa for each of five taxonomic levels. Relative abundance frequencies are plotted for taxa levels from both the non-rarefied and the rarefied datasets.

**Figure S2 - Heatmaps showing the genes involved in any function that were found enriched by IPA gene set analysis.** Taxon-specific analysis on genes in the QTL regions associated with relative abundance of phylum Firmicutes **(A)**, genus *Staphylococcus* **(B)**, family Ruminococcaceae **(C)**, class Mollicutes **(D)**, and order Bacteroidales **(E)**. We only show annotations with a False Discovery rate under 3% after multiple-hypothesis correction. Filled-in cells indicate that the gene listed at the top of that column is annotated with the function or disease of that row. Only genes in the gene set of interest are shown, these charts do not display all gene members of each pathway.

**Figure S3**- **Correlation plot between non-rarefied and rarefied taxa.** Heatmap depicting the Pearson correlations between the relative common taxa relative abundances in non-rarefied (NonR) and rarefied (R) data, revealing that the same taxa from both non-rarefied and rarefied datasets always group closer together than with other taxa, followed by taxa belonging to the same clade.

**Figure S4 - Proportion of variance estimates for kinship and cage for all taxa in rarefied data.** Proportion of variance estimates for kinship (green), cage effects (orange), and unexplained residual effects (blue) for each taxon. The kinship proportion of variance is an estimate of narrow sense heritability. Heritability percentages are shown on the left. Heritability standard error values are shown with black horizontal lines. Designations p_, c_, o_, f_, and g_ are for phylum, class, order, family, and genus, respectively. When results are identical across taxa in the same phylogenetic branch, only the lowest (most specific) taxa are shown and the rest are shaded out. Heritability significance is marked with one plus (**+**, RLTR *p*-value < 0.05) and BH FDR is shown in parentheses next to heritability percentages. Taxa marked with a red asterisk have statistically suggestive QTL (⭑, adj. *p*-value < 0.1). Complete table of heritability results, including non-rarefied data, can be found in **Table S3**.

**Figure S5 - Comparison of heritability estimates between non-rarefied and rarefied taxa**. Circles with purple fill correspond to non-rarefied taxa with statistically significant heritabilities. Circles with green outlines correspond to rarefied taxa with statistically significant heritabilities. Standard errors are shown as horizontal blue lines for non-rarefied taxa and vertical orange lines for rarefied taxa.

**Table S1 - Relative abundance of OTUs.** Relative microbial abundance at the OTU level for each DO mouse in non-rarefied data (**A**) and rarefied data (**B**).

**Table S2 - Microbial relative abundance summarized at five levels of taxonomy.** Relative microbial abundance summarized at five levels of taxonomy (phylum, class, order, family, and genus) for each DO mouse in non-rarefied data (**A**) and rarefied data (**B**).

**Table S3**- **Heritability results at five taxonomic levels.** Complete heritability measurements (*h^2^*) as well as their respective *p*-values, adjusted *p*-values, and standard errors for all tested taxonomies at the five taxonomic levels from the non-rarefied (**A**) and rarefied (**B**) datasets.

**Table S4**- **QTL results at five taxonomic levels.** QTL regions and their respective *p*-values, permutation *p*-values (when applicable), and genes found within the QTL interval at the five taxonomic levels from the non-rarefied (**A**) and rarefied (**B**) datasets.

**Table S5 - Genes within QTL regions with suggestive permutation***p***-value.** Detailed annotations for all genes found within QTL regions with a permutation *p*-value <0.1 at the five taxonomic levels.

**Table S6**- **QTL results at OTU level in non-rarefied dataset.** QTL regions and their respective *p*-values, permutation *p*-values (when applicable), and genes found within the QTL interval at the OTU level from the non-rarefied dataset.

**Table S7**- **Comparison of heritabilities and QTL with other studies.** Comparison of taxa heritabilities and QTL from our analyses with other studies across mouse, human, and pig studies (**A**). Information on source studies for heritability values in (**B**) and for QTL/GWAS in (**C**). Full human to mouse synteny mapping results for human studies in (**D**).

**Table S8 - Gene network relationships from Figure 8.** Annotated relationships between the genes in network 1 (**A**), network 2 (**B**), and network 3 (**C**) from **Figure 8**.

**Table S9 - Functional annotation table from IPA analysis.** Detailed functional annotations from cumulative gene set enrichment analysis using IPA on 1423 genes associated with non-rarefied microbiome abundance.

