## Supplementary material for "High-resolution QTL mapping with Diversity Outbred mice identifies genetic variants that impact gut microbiome composition": Fig S1

Phylum Non-Rarefied

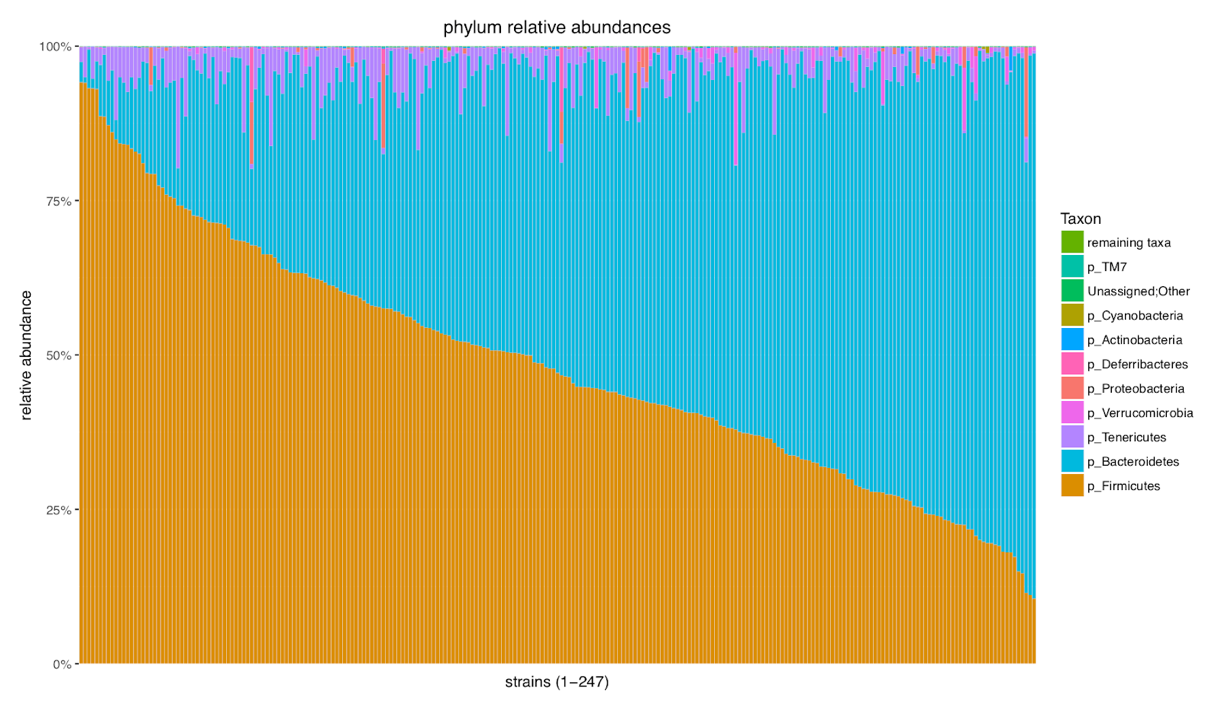

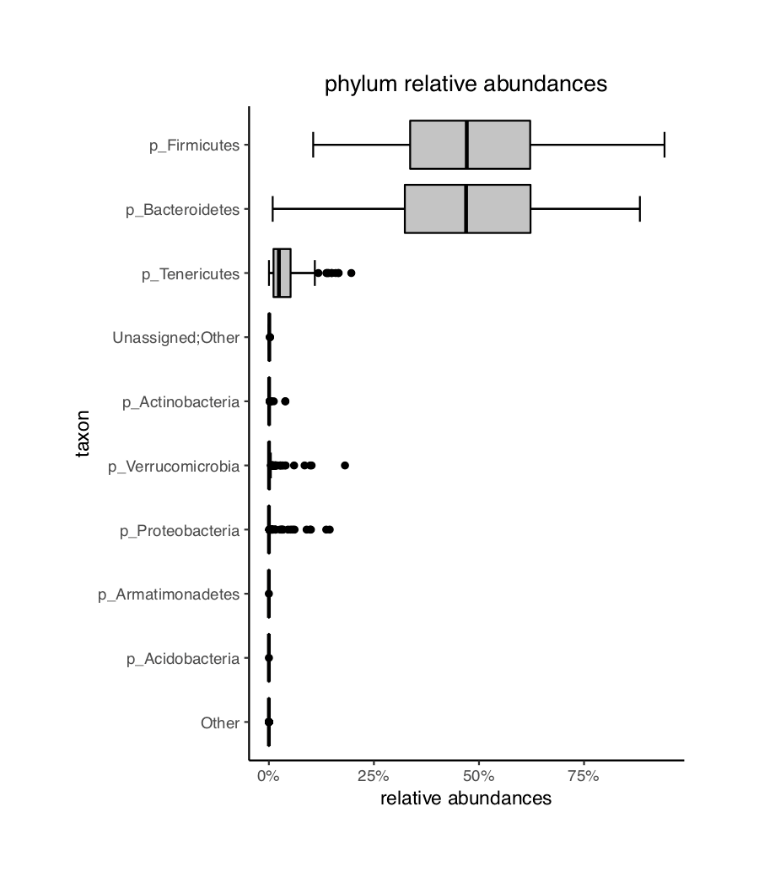

Phylum Rarefied
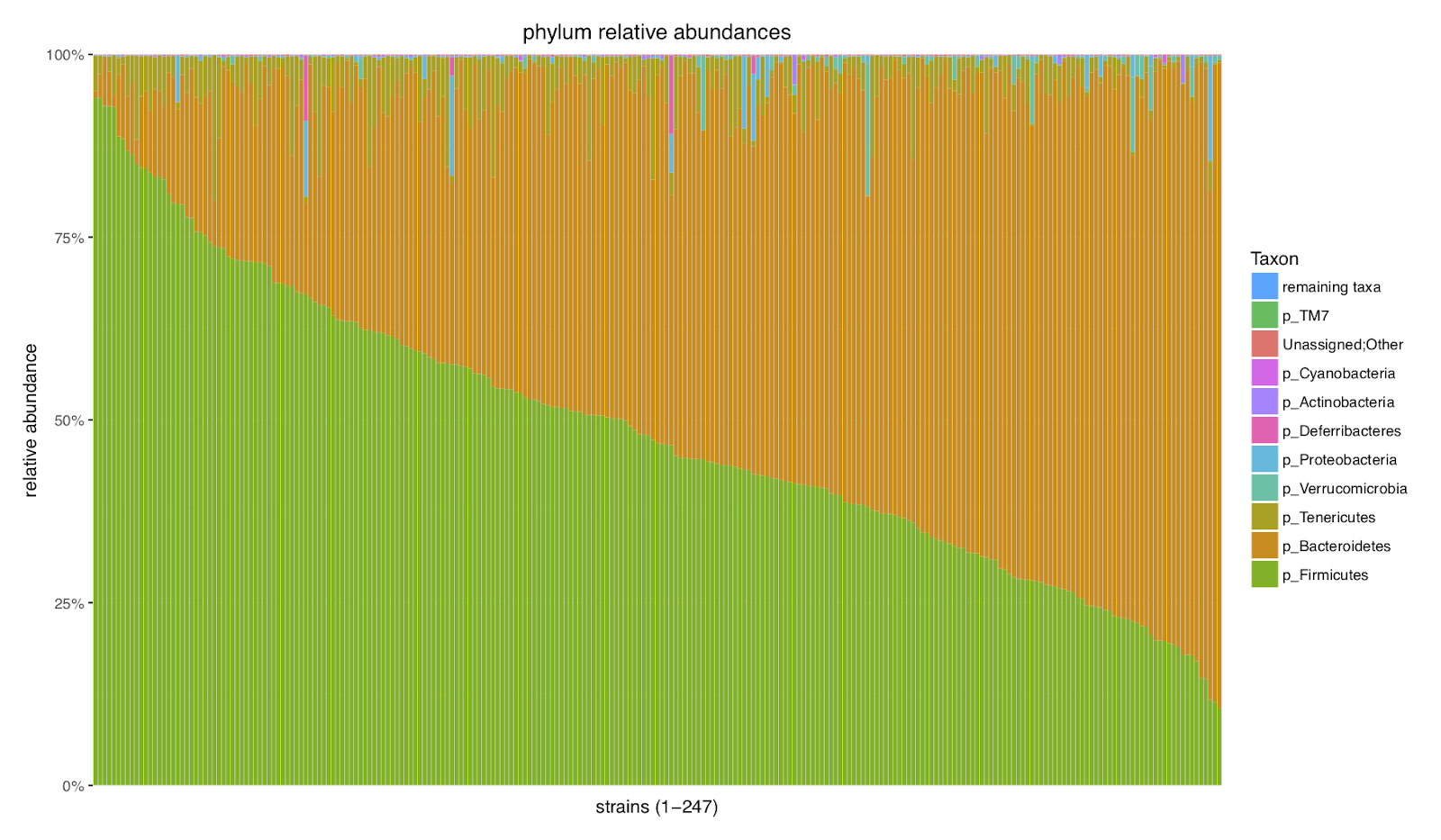

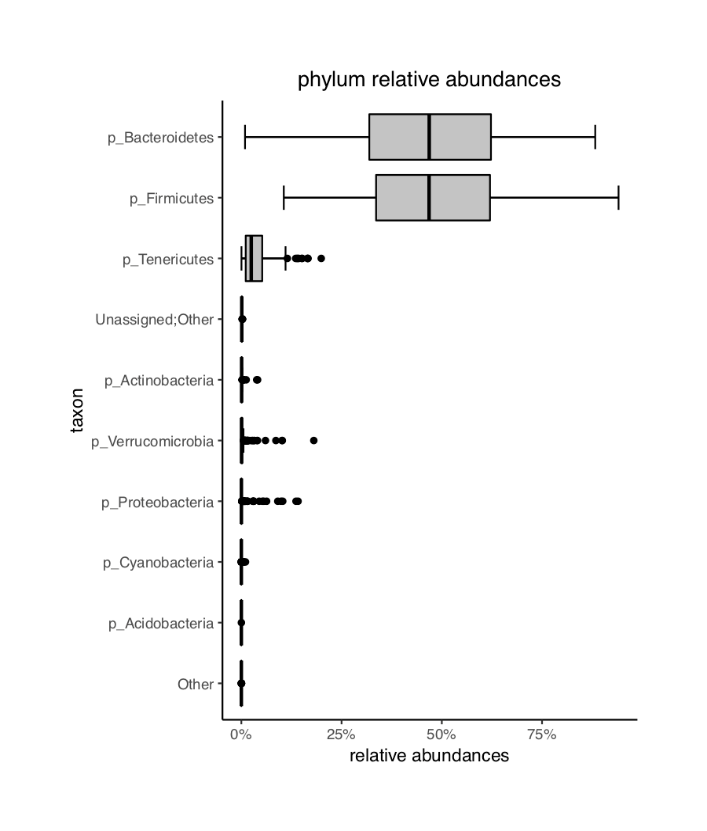

Class NonR
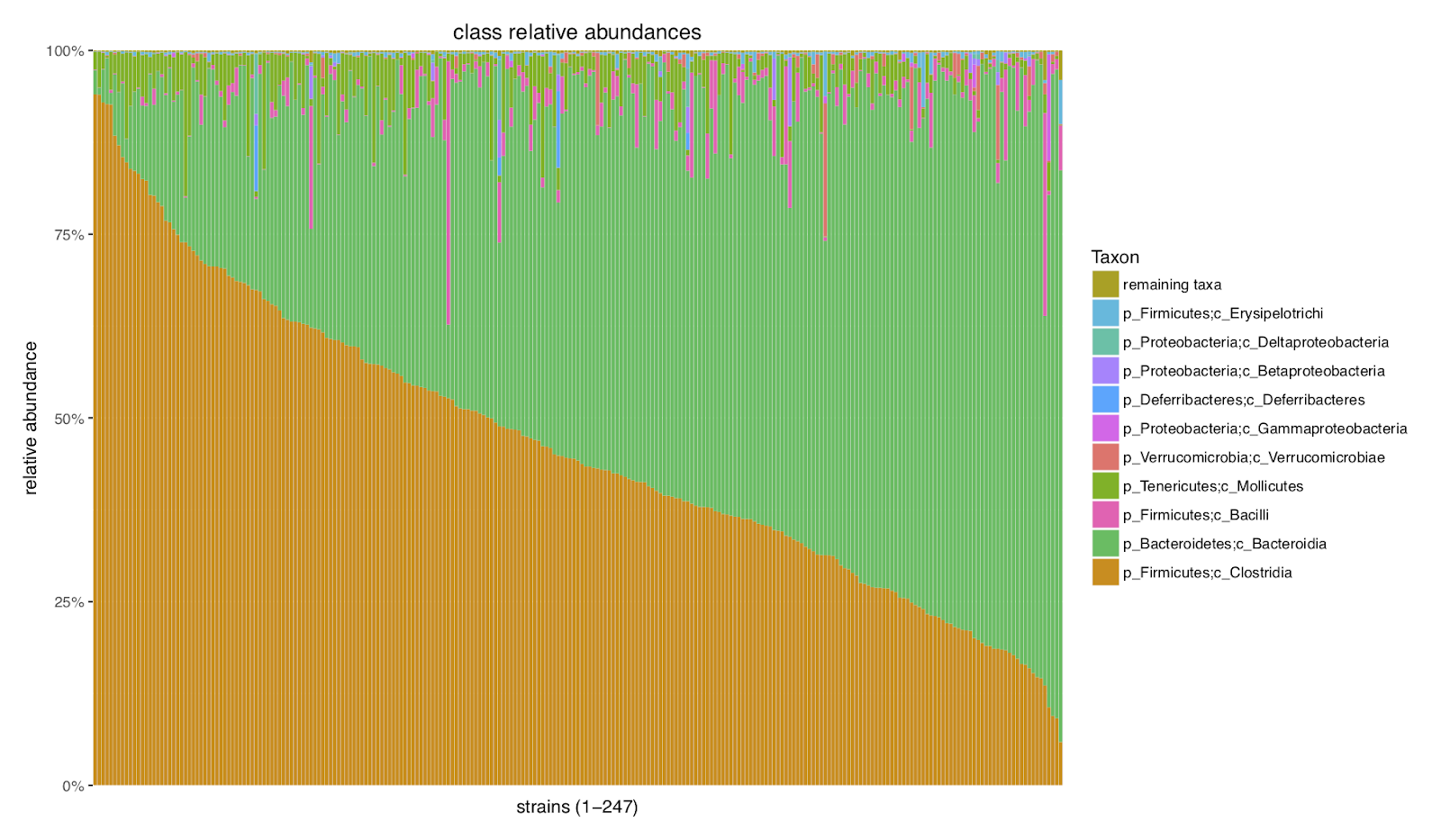

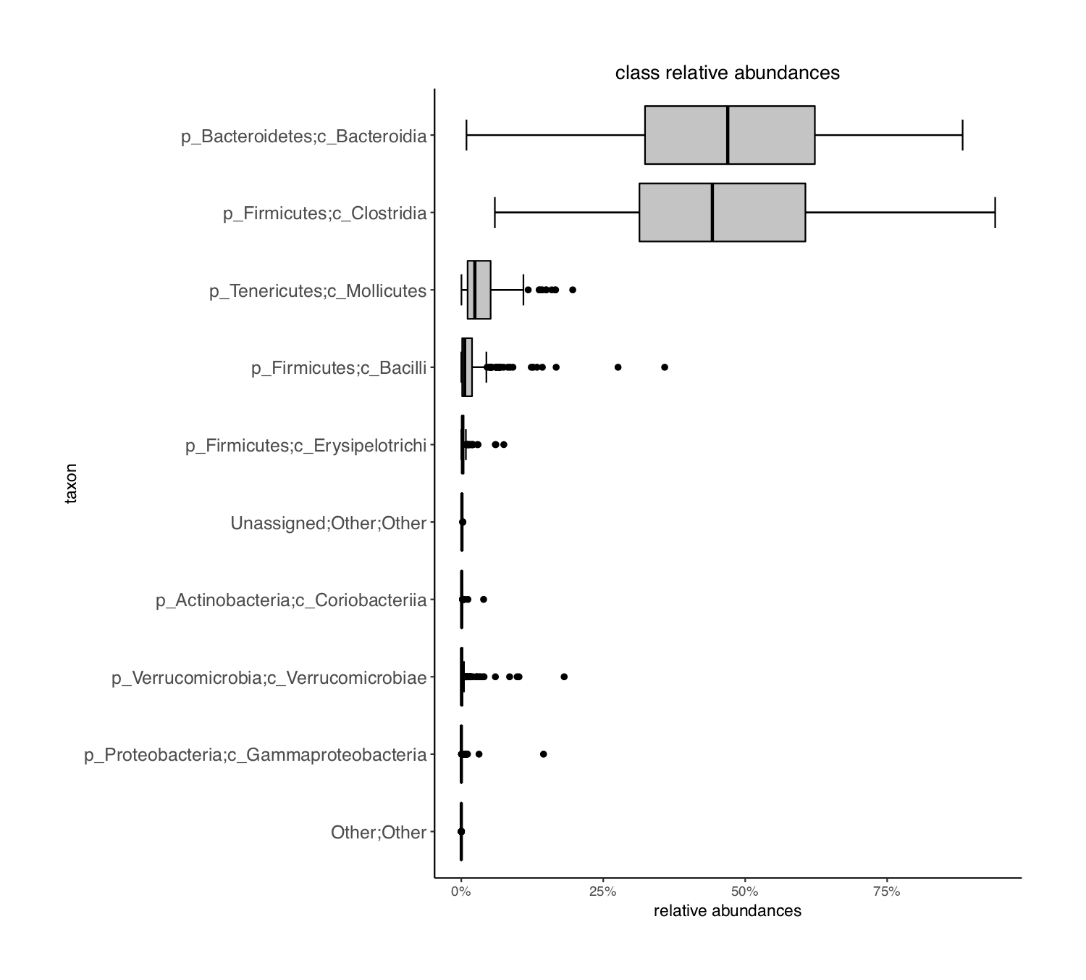

Class R

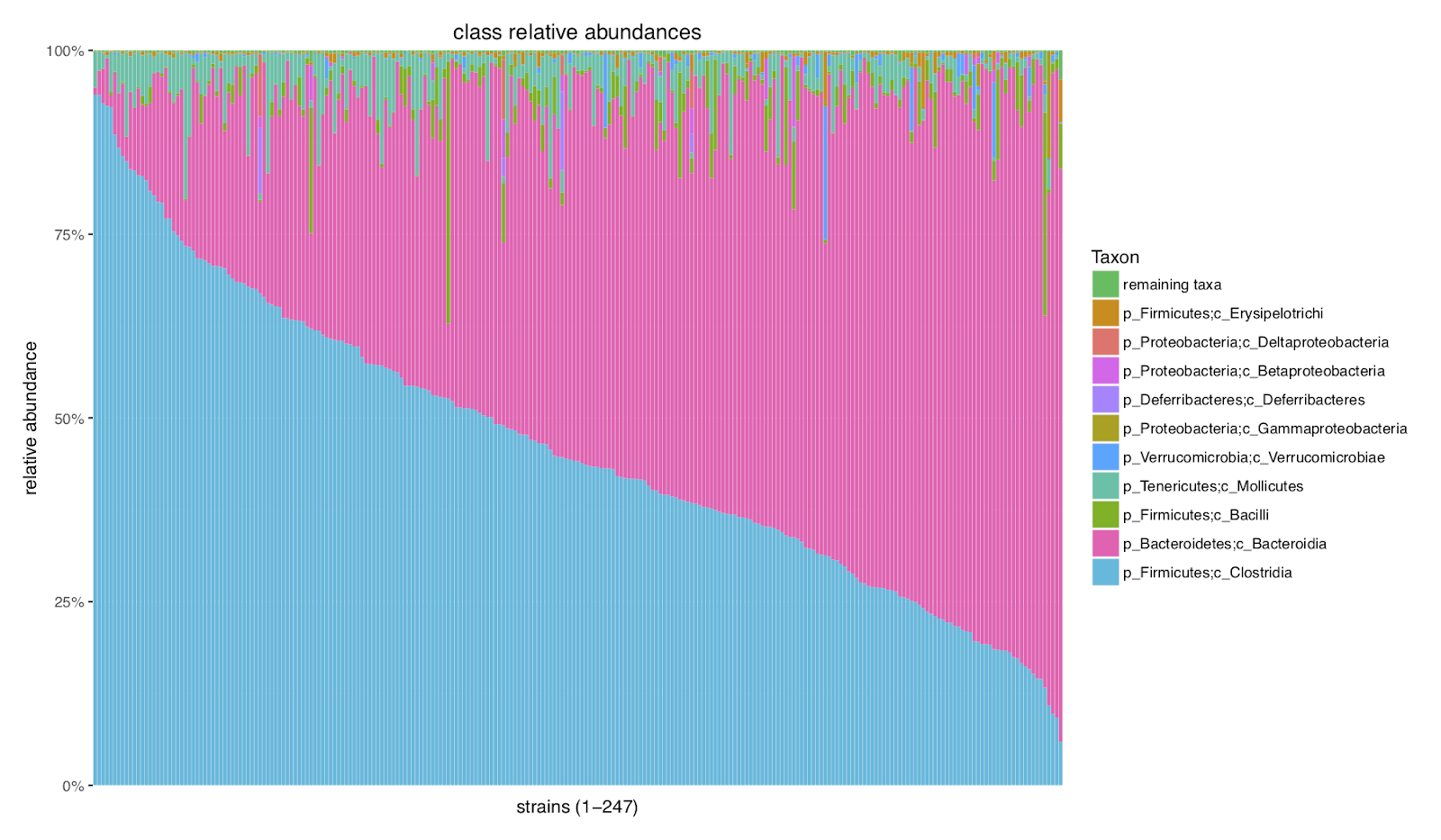

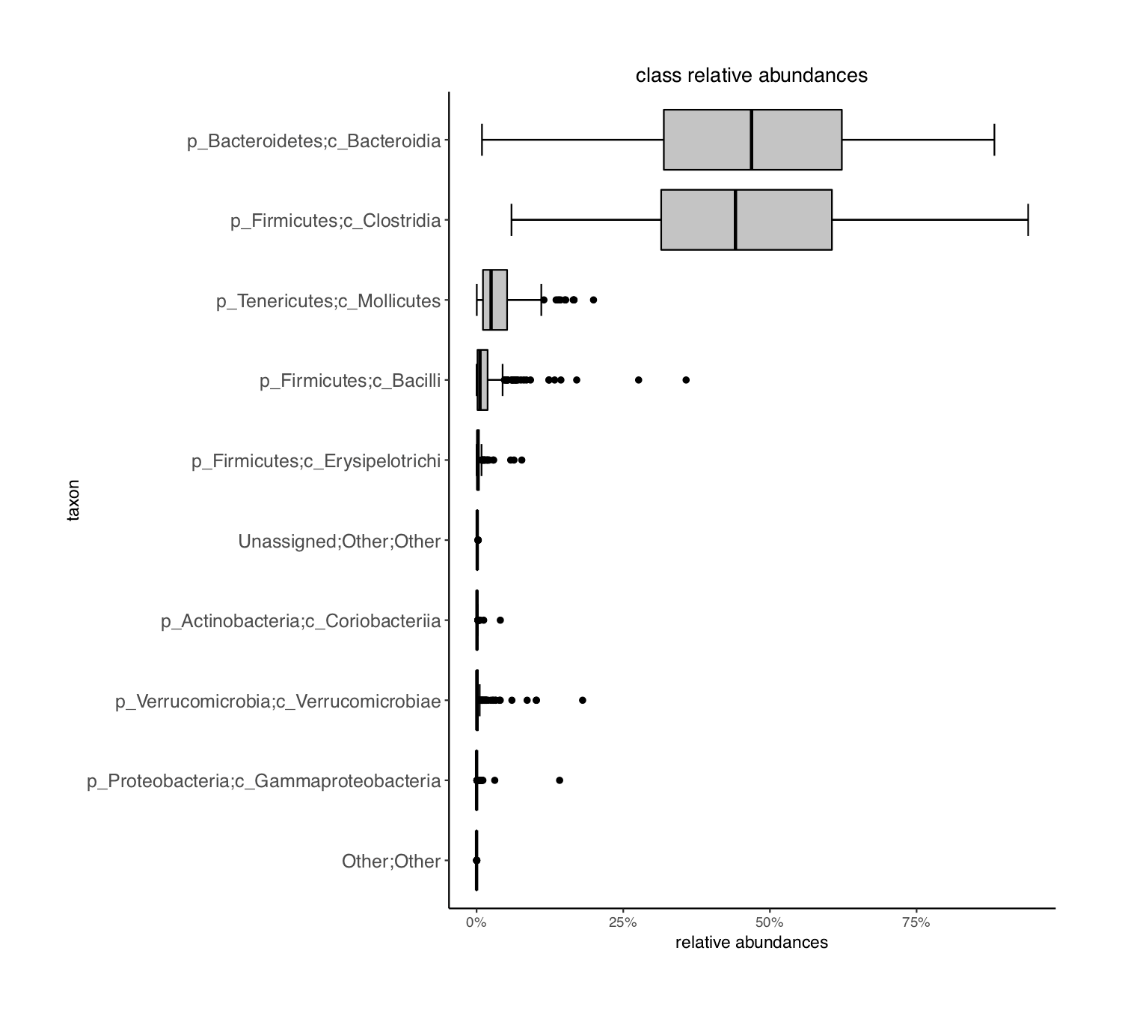

Order NonR

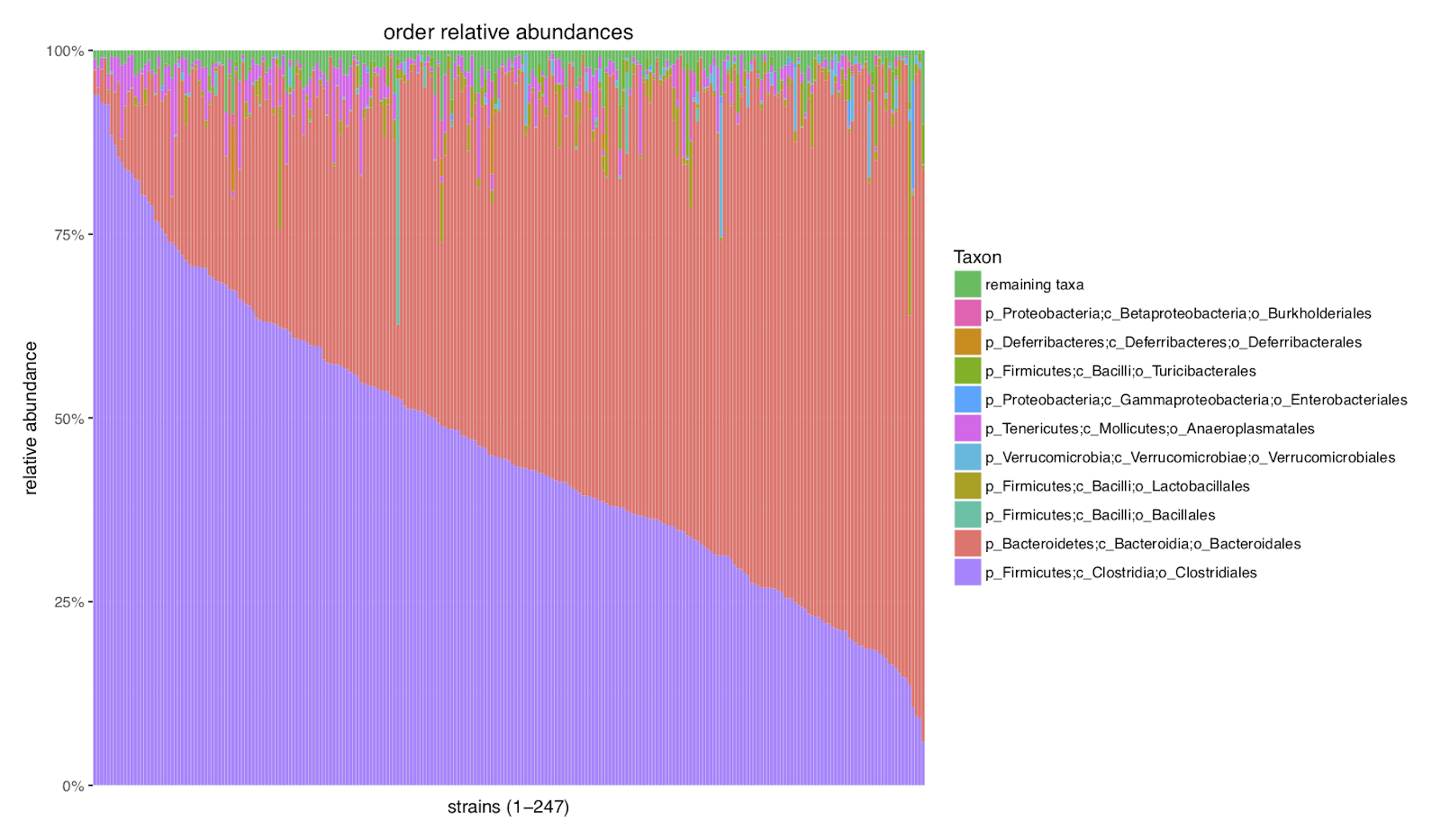

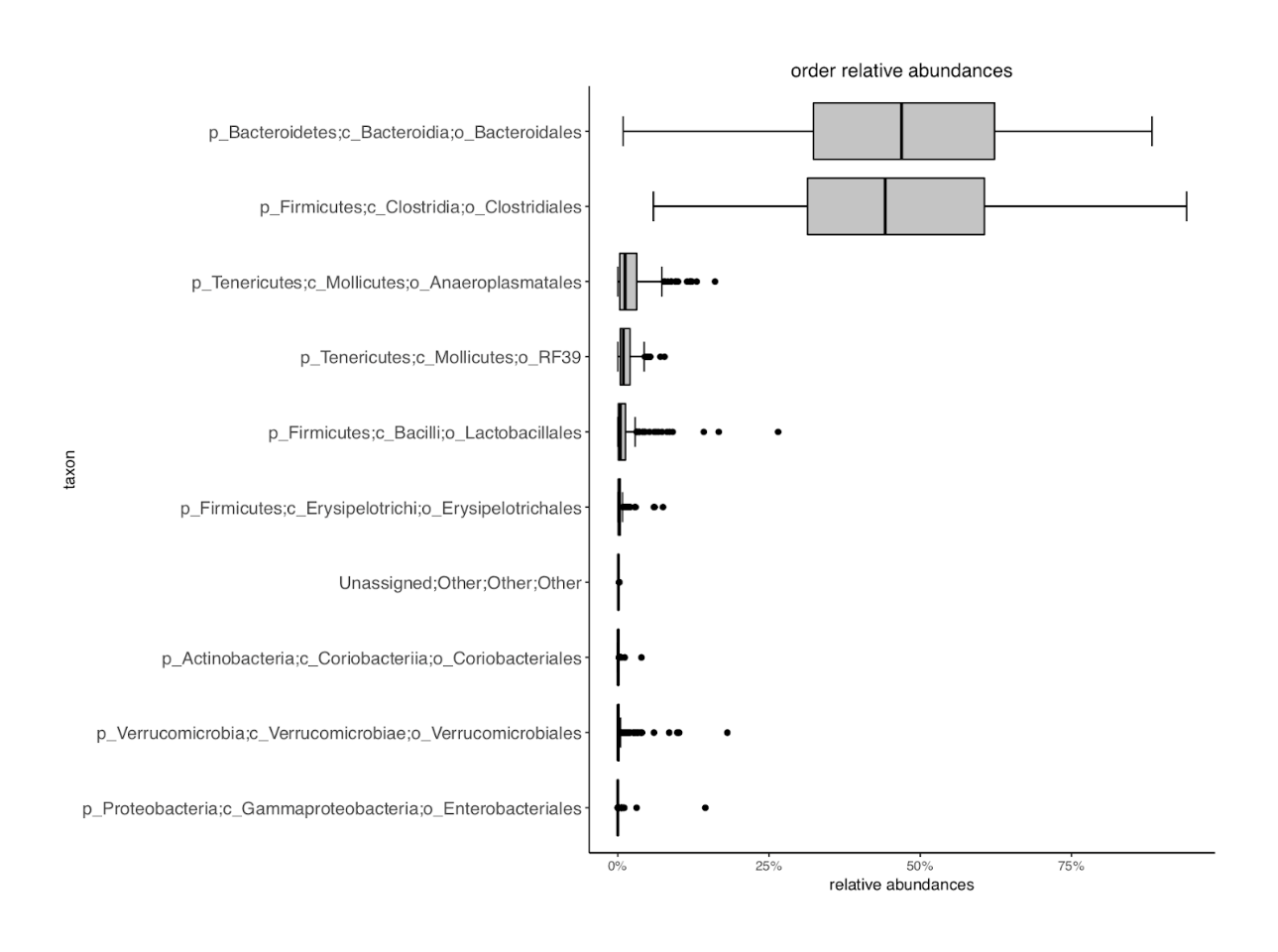

Order R

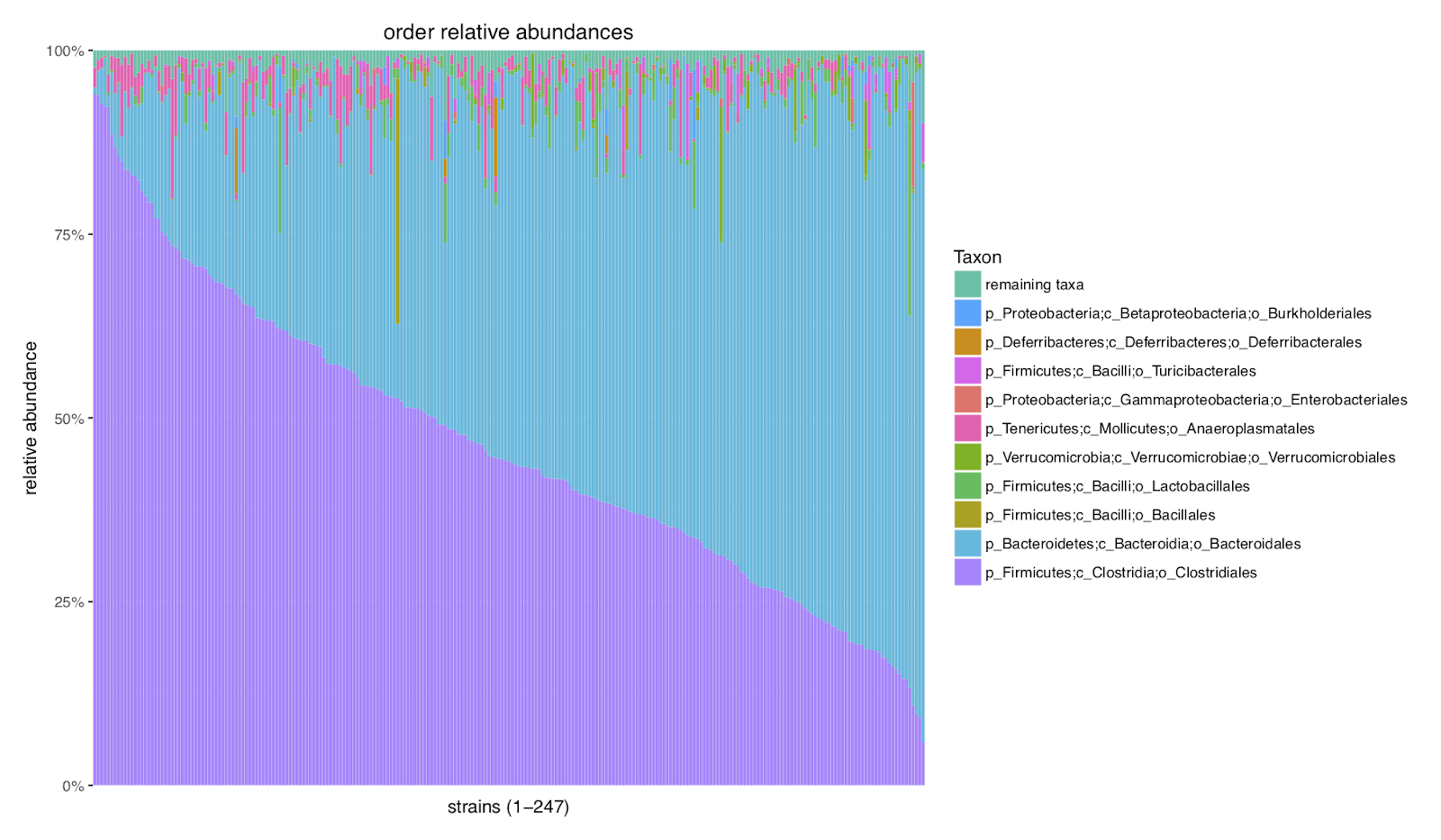

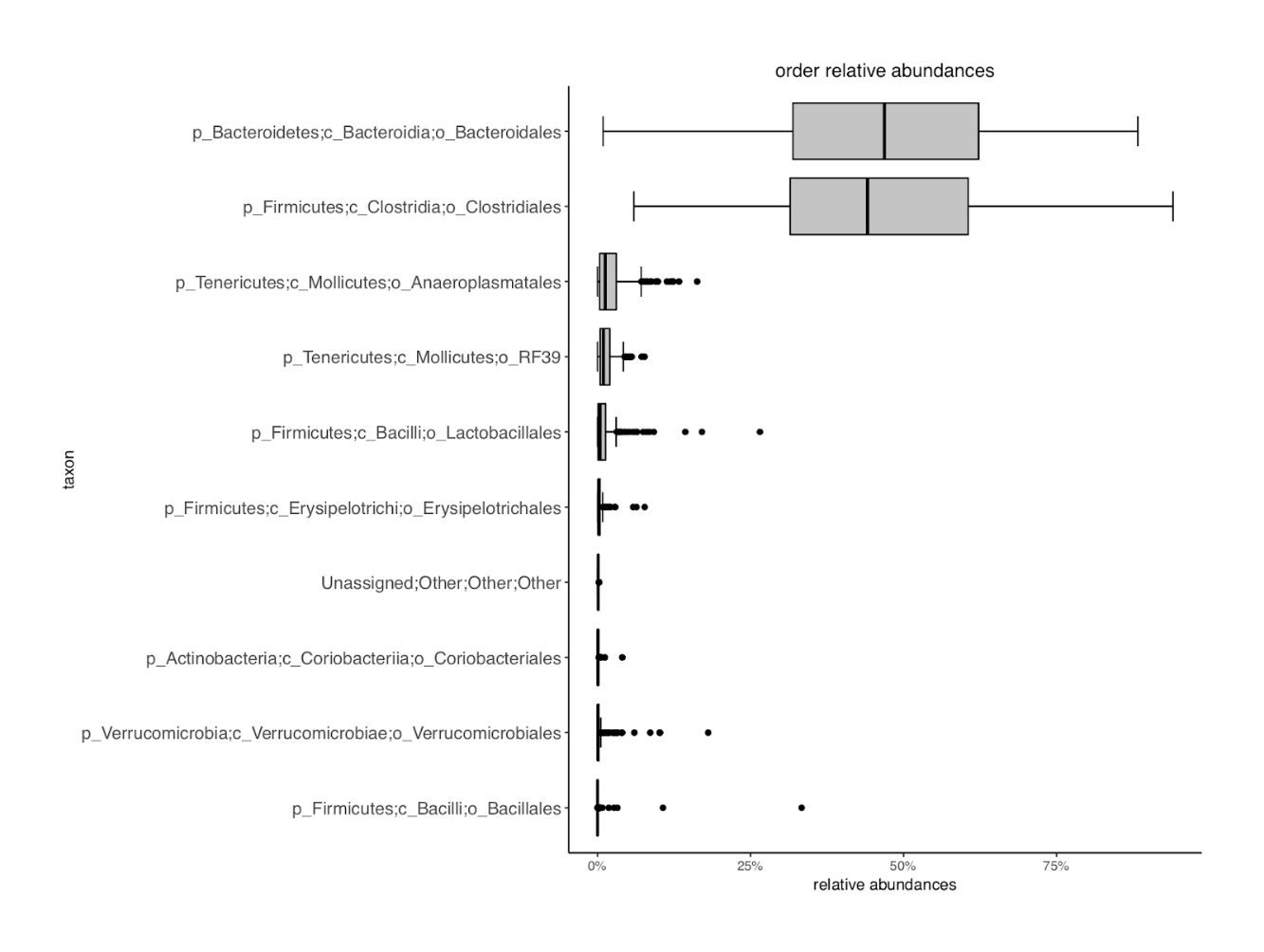

Family NonR

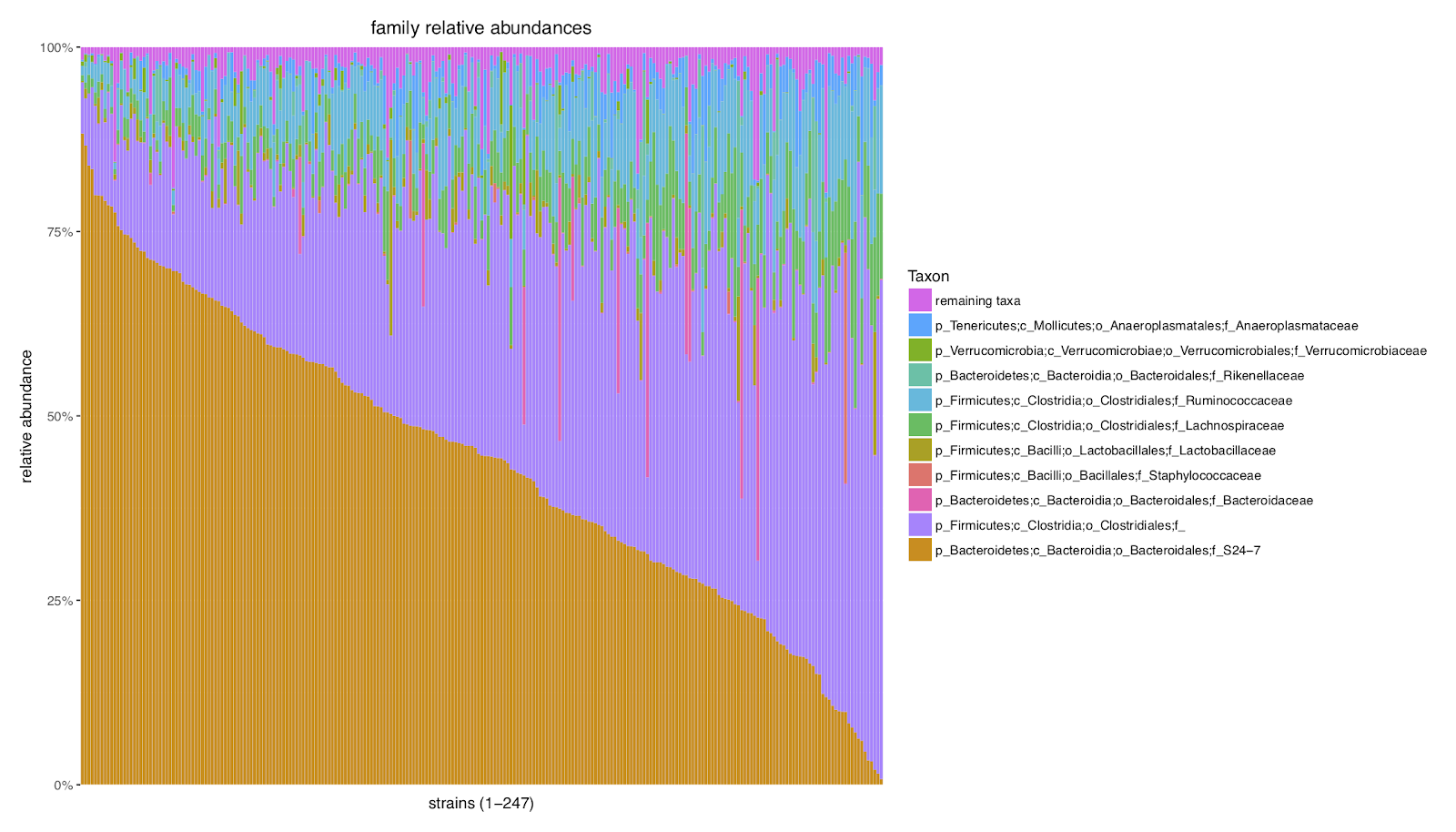

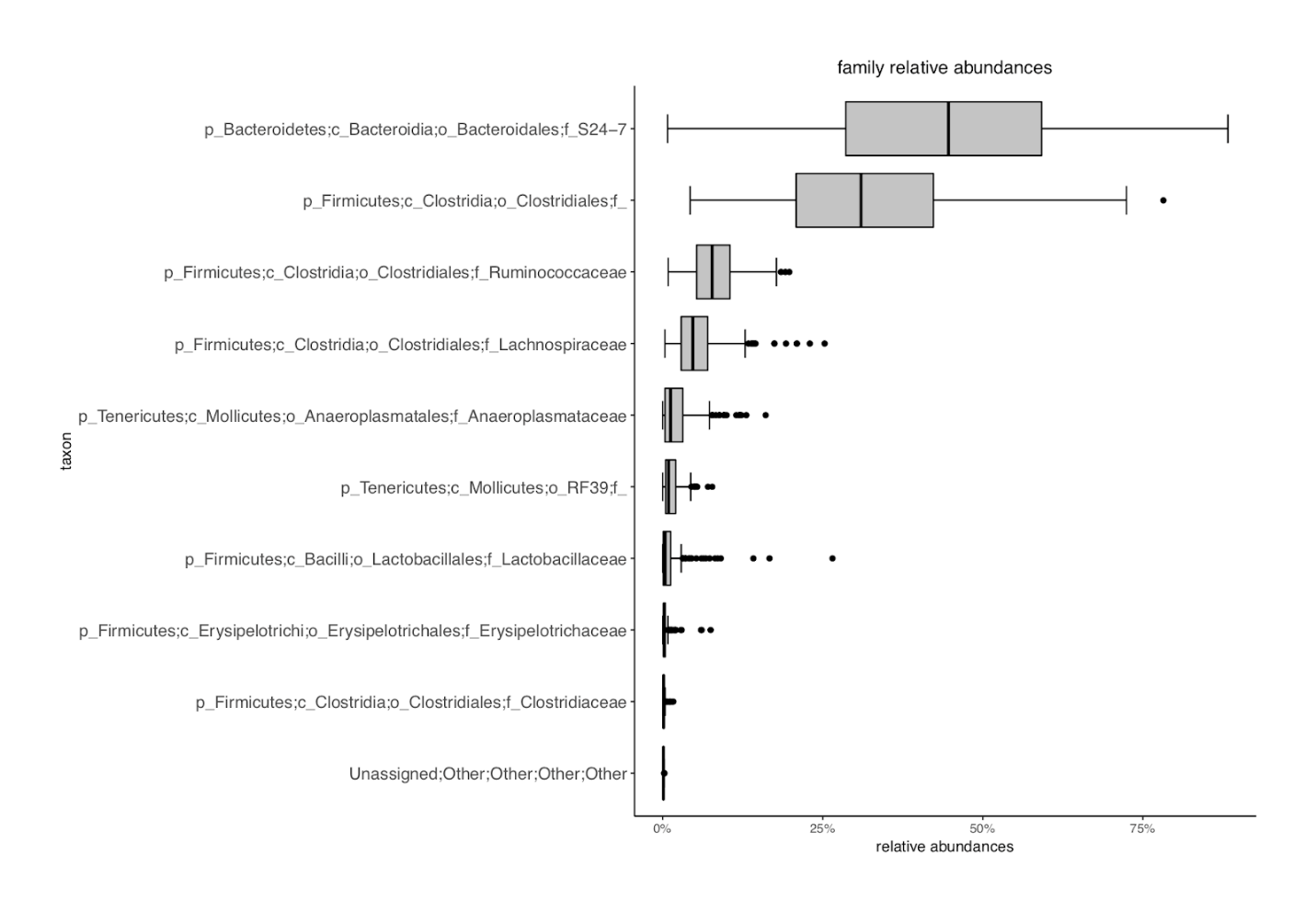

Family R
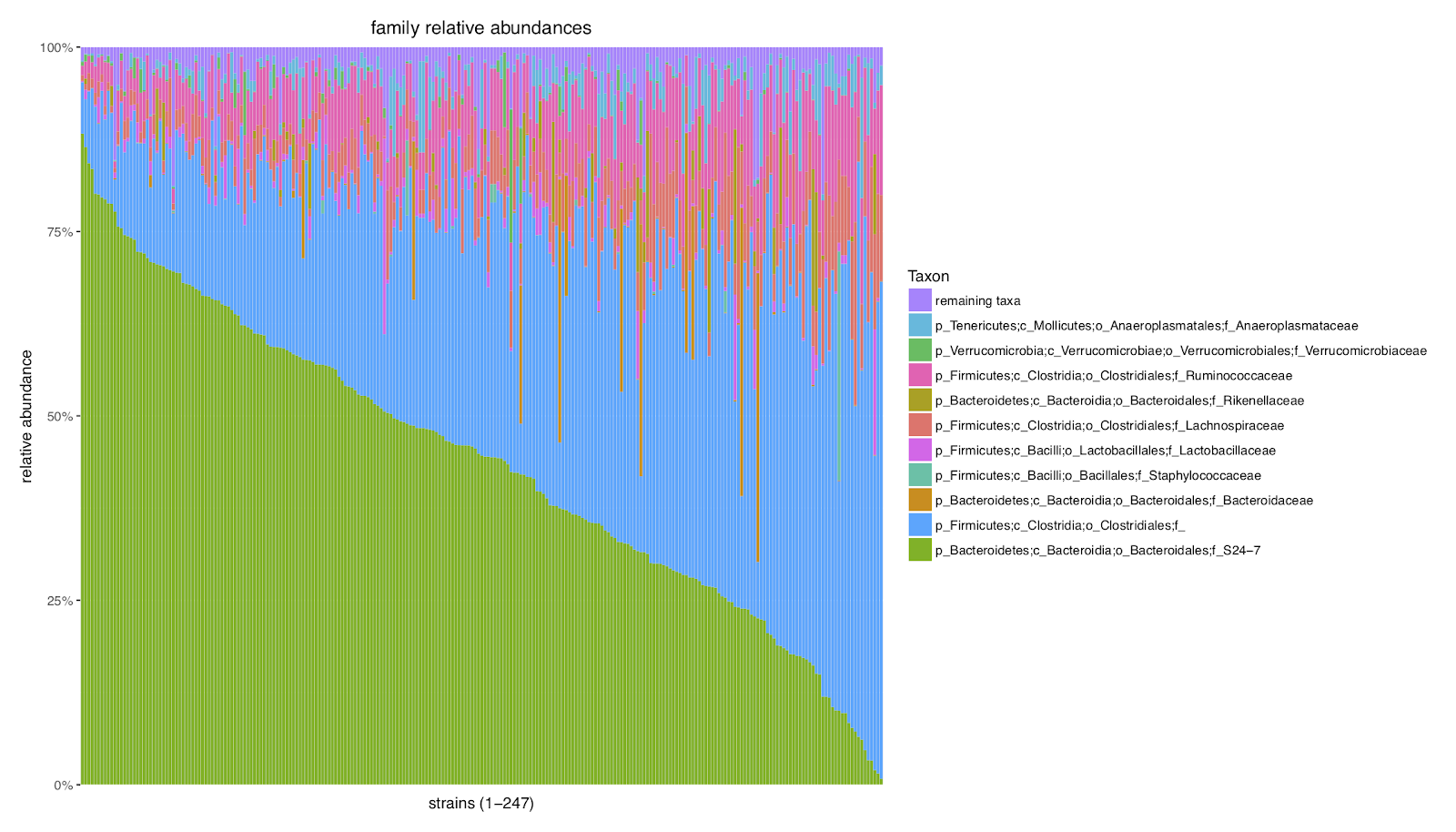

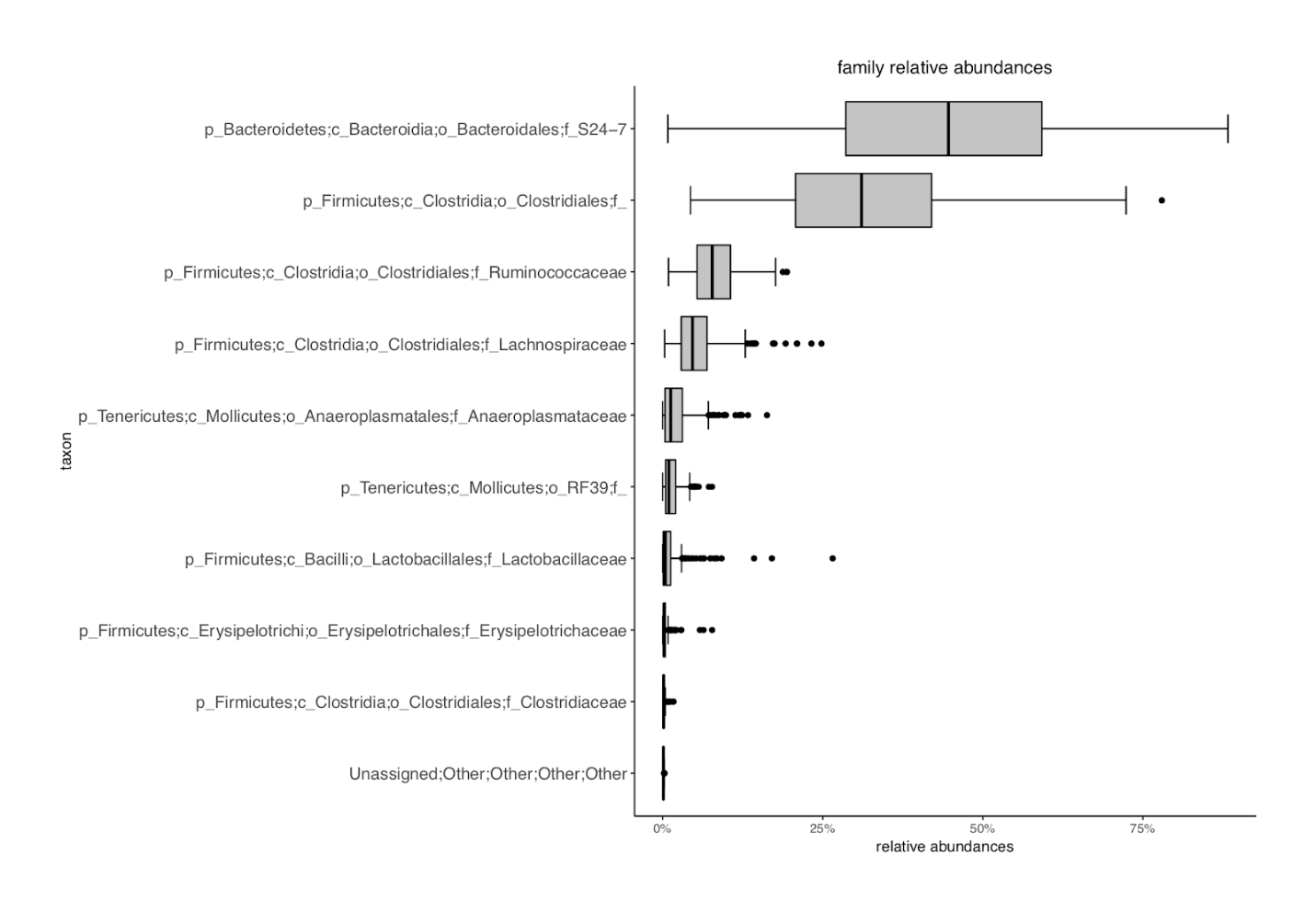

Genus NonR
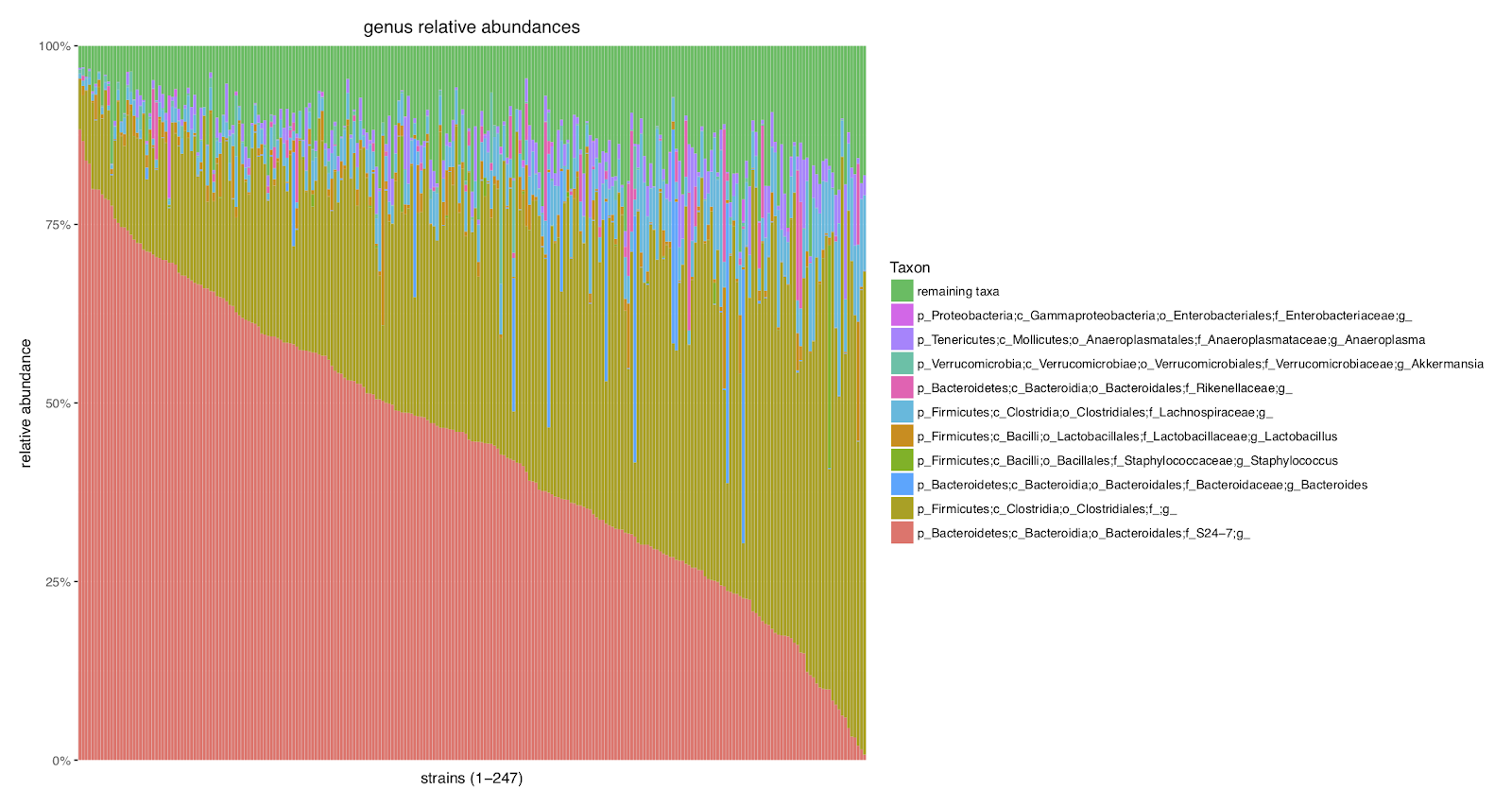

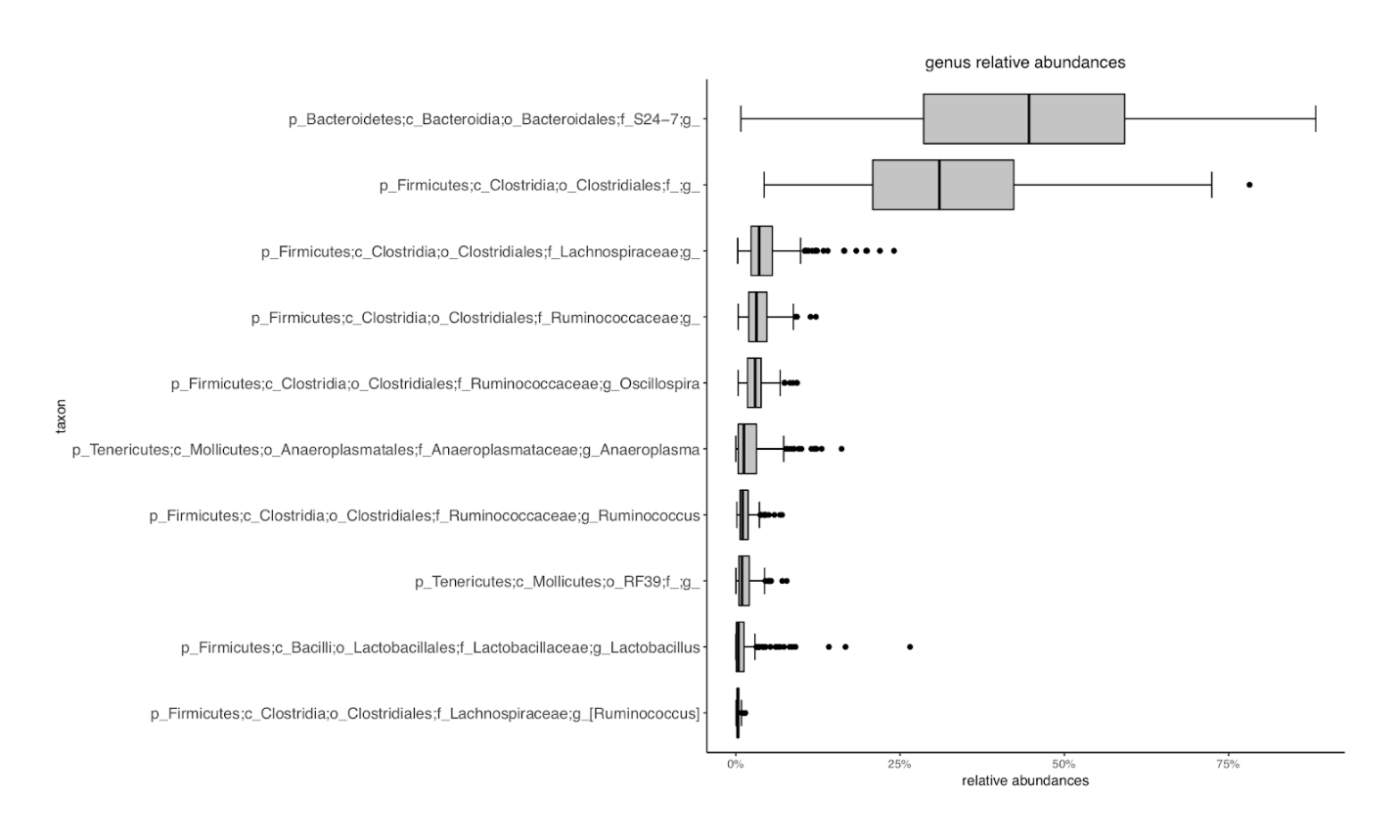

Genus R
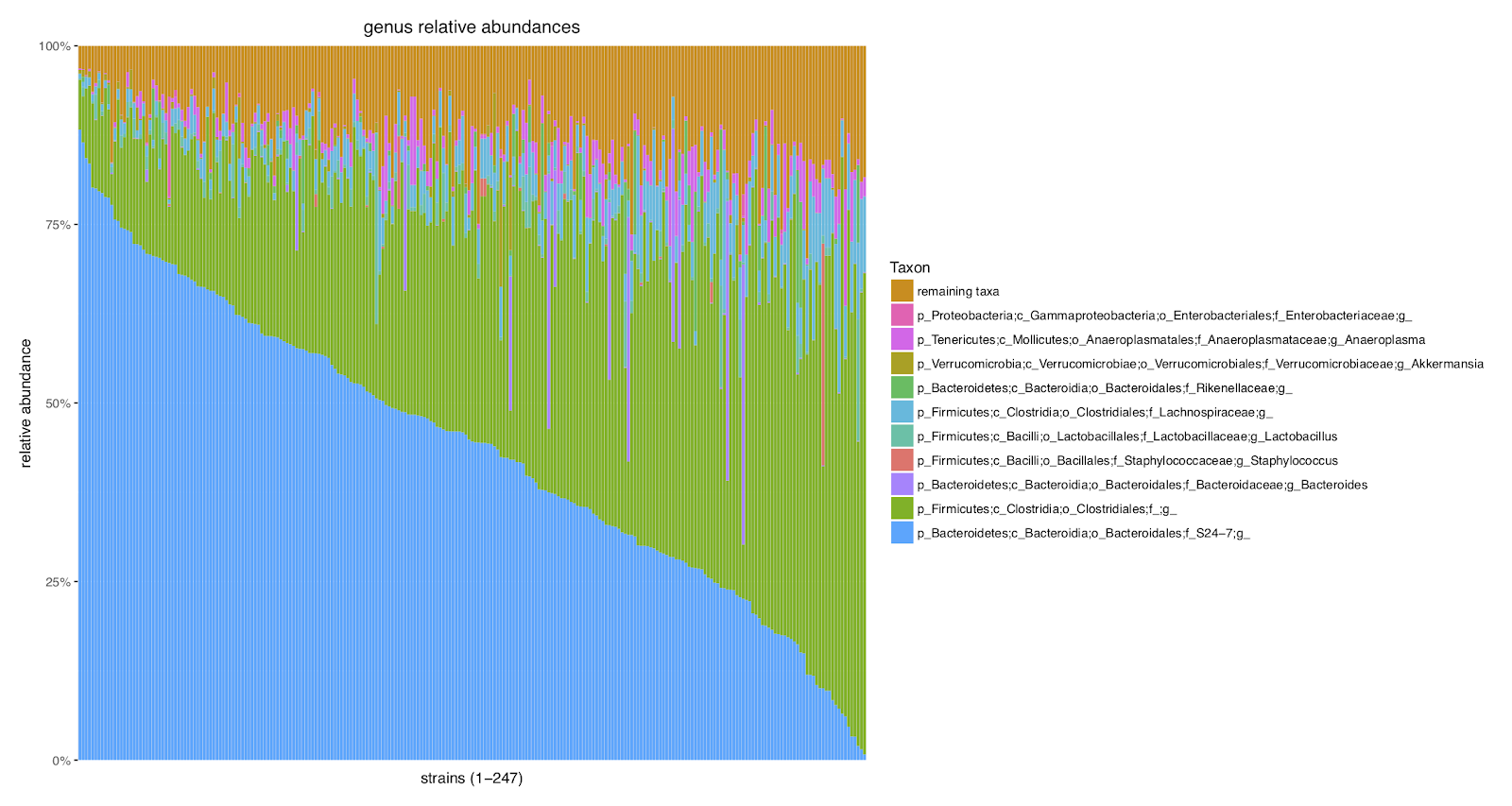

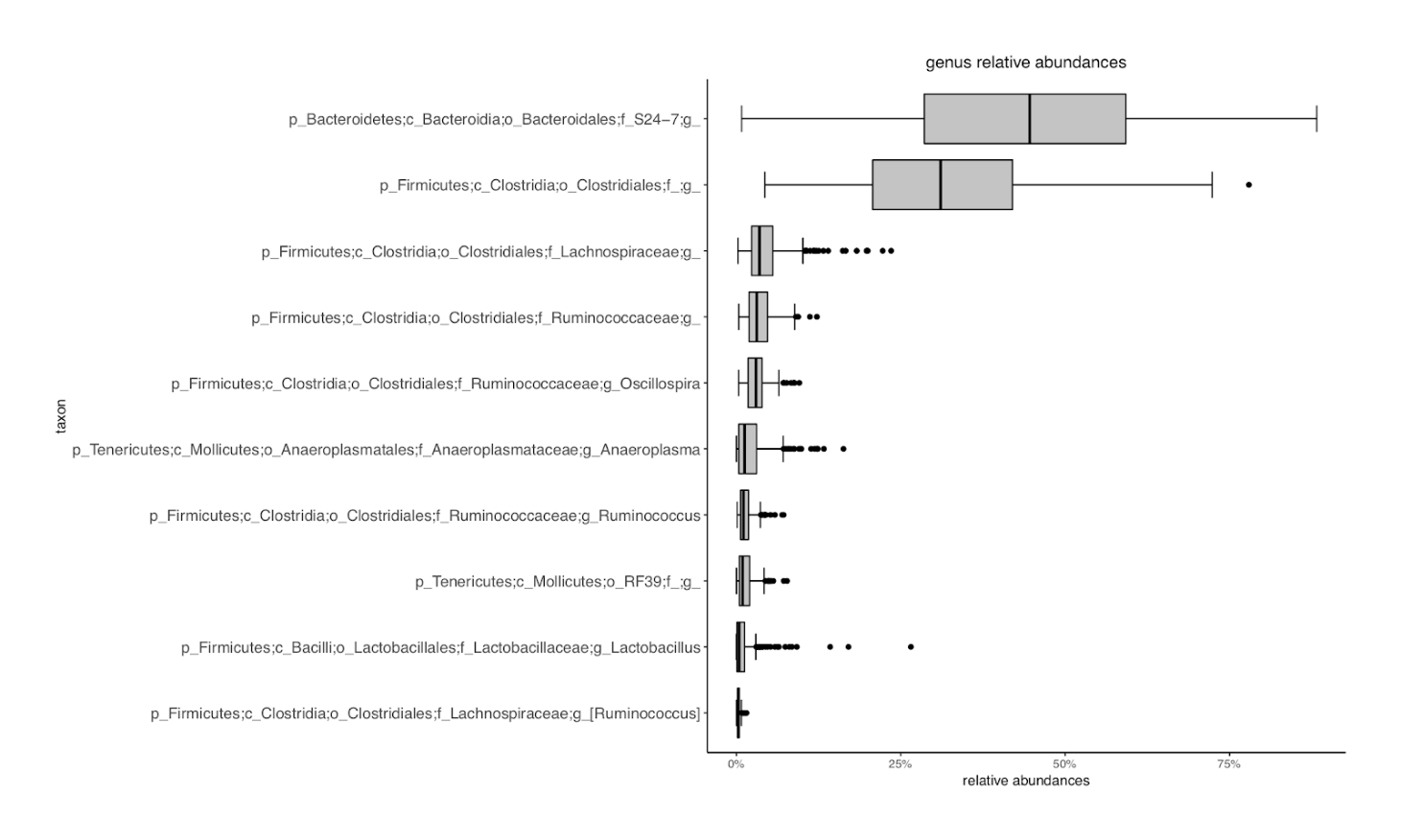

**Figure S1**. **Taxa relative abundance frequencies**. Stacked bar plots and box plots depicting relative abundance frequencies of the top ten most abundant taxa for each of five taxonomic levels. Relative abundance frequencies are plotted for taxa levels from both the non-rarefied and the rarefied datasets.
