## Supplementary material for "High-resolution QTL mapping with Diversity Outbred mice identifies genetic variants that impact gut microbiome composition": Fig S2

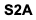

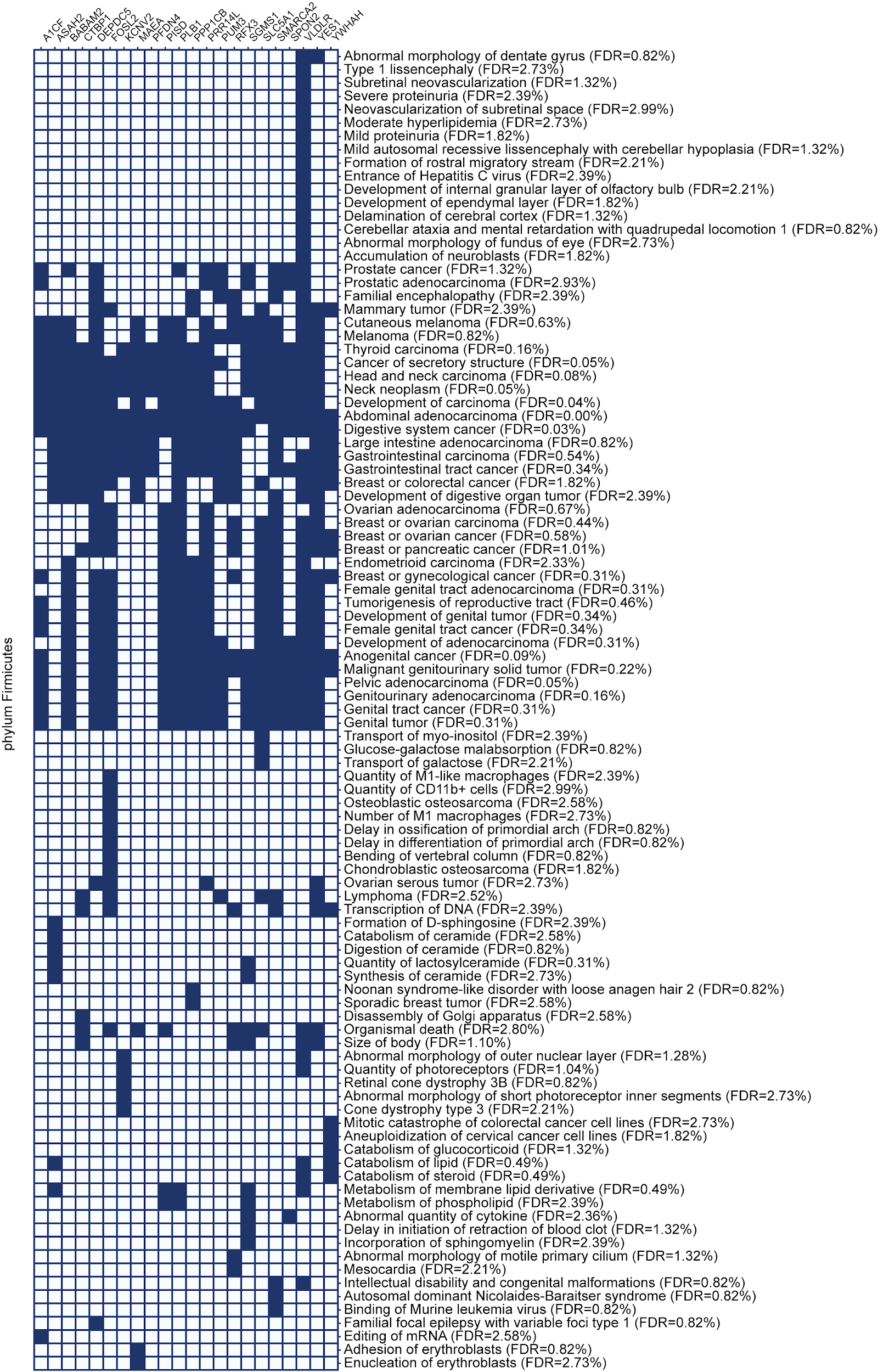

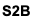

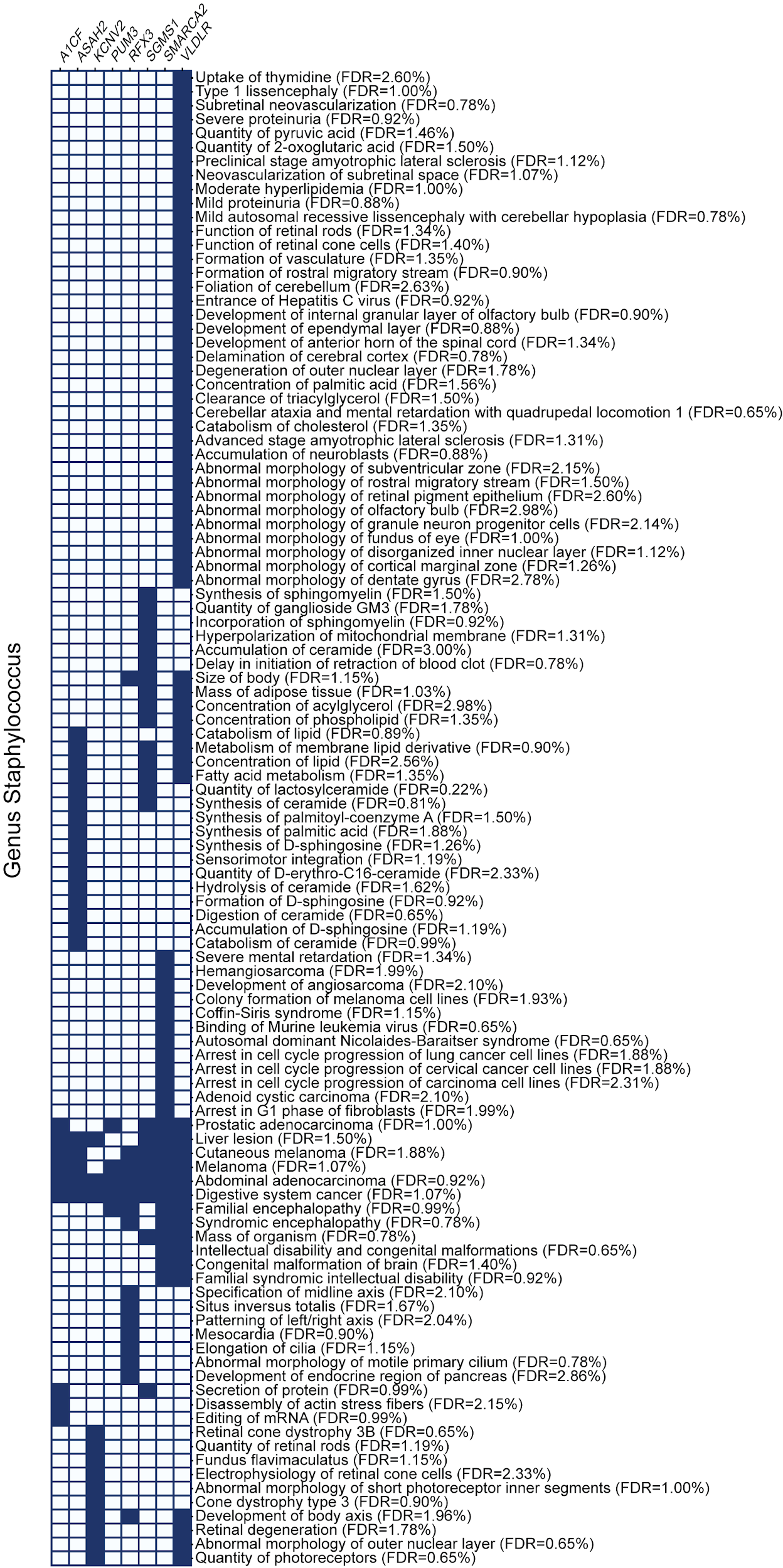

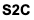

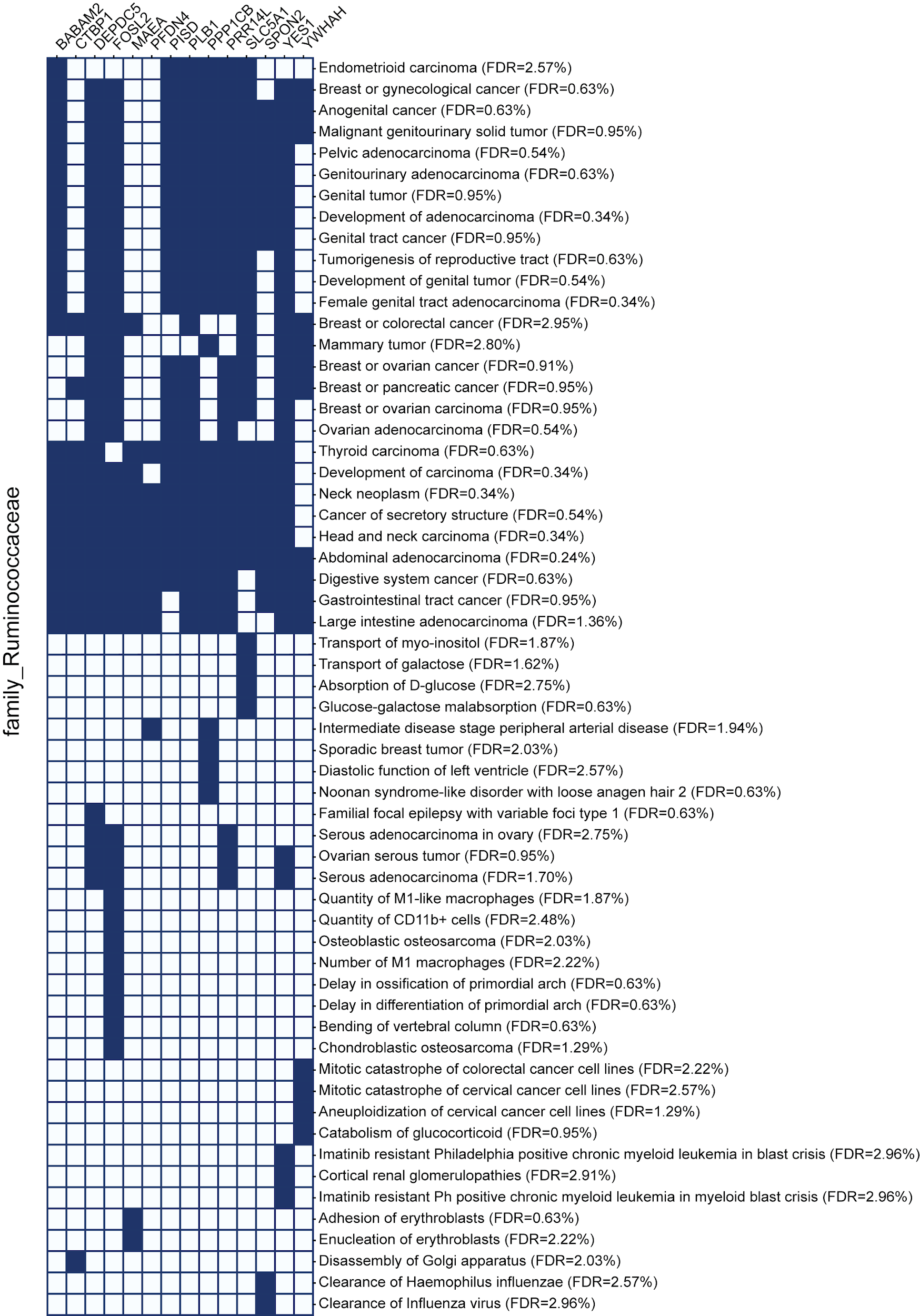

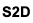

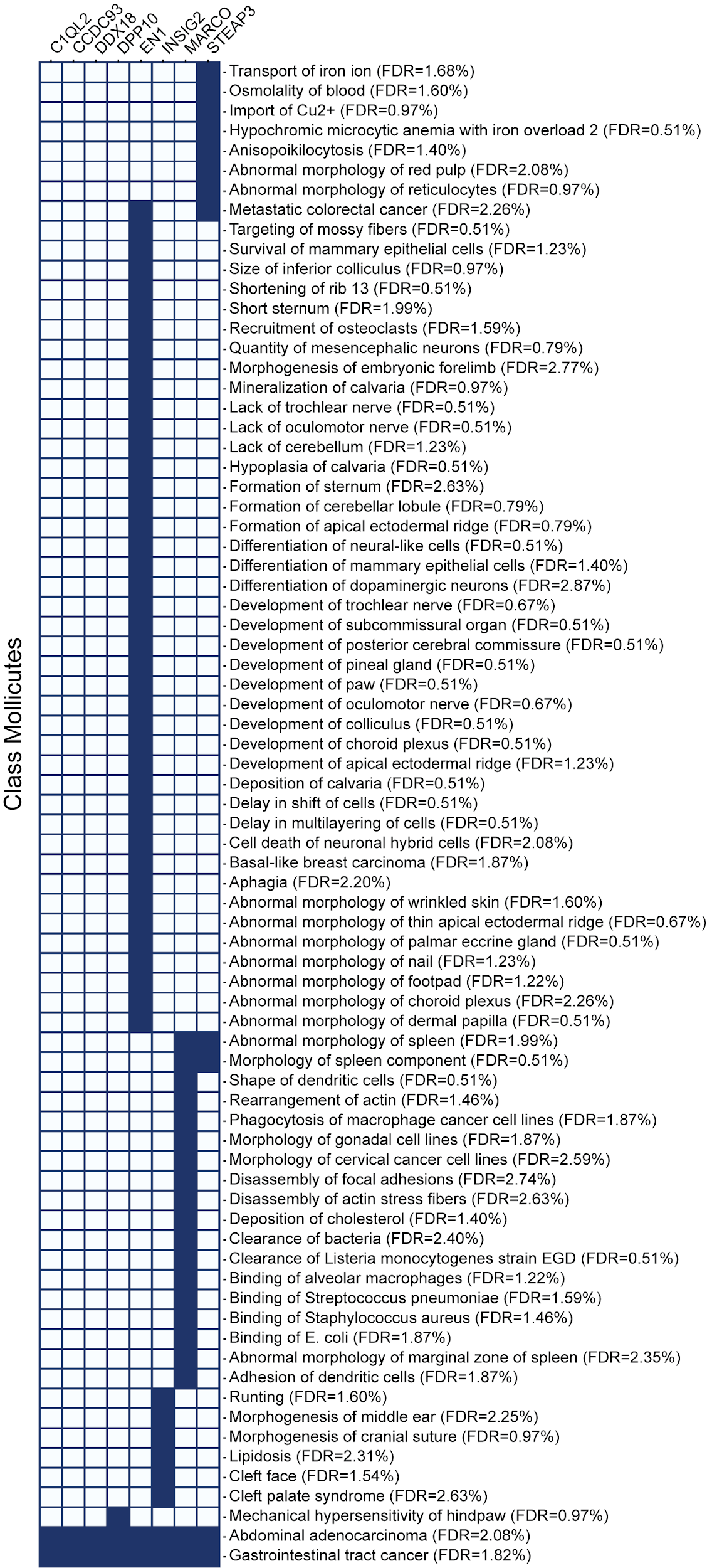

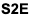

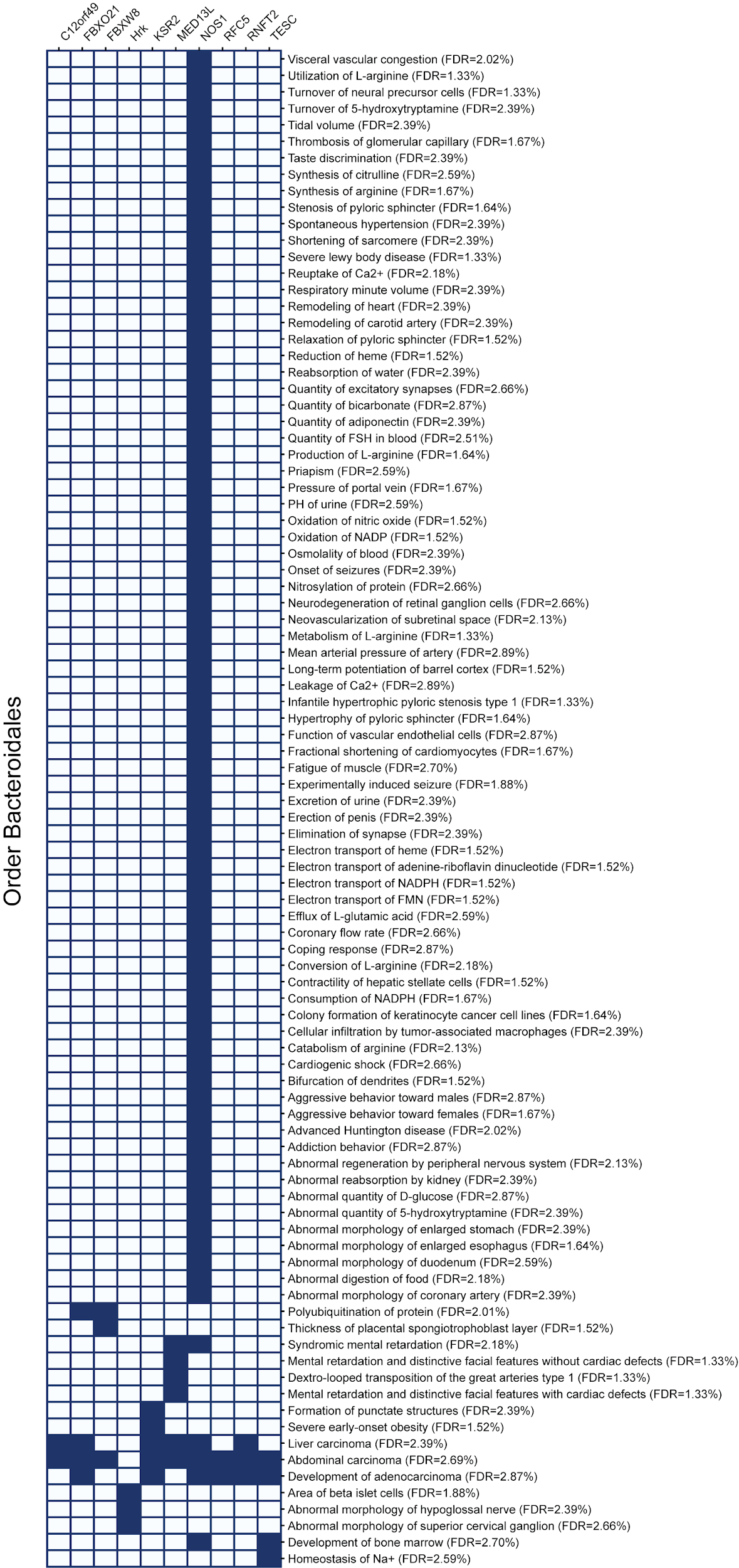

**Figure S2**. **Heatmaps showing the genes involved in any function that were found enriched by IPA gene set analysis.** Taxon-specific analysis on genes in the QTL regions associated with relative abundance of phylum Firmicutes **(A)**, genus *Staphylococcus* **(B)**, family Ruminococcaceae **(C)**, class Mollicutes **(D)**, and order Bacteroidales **(E)**. We only show annotations with a False Discovery rate under 3% after multiple-hypothesis correction. Filled-in cells indicate that the gene listed at the top of that column is annotated with the function or disease of that row. Only genes in the gene set of interest are shown, these charts do not display all gene members of each pathway.
