## Supplementary material for "High-resolution QTL mapping with Diversity Outbred mice identifies genetic variants that impact gut microbiome composition": Fig S3

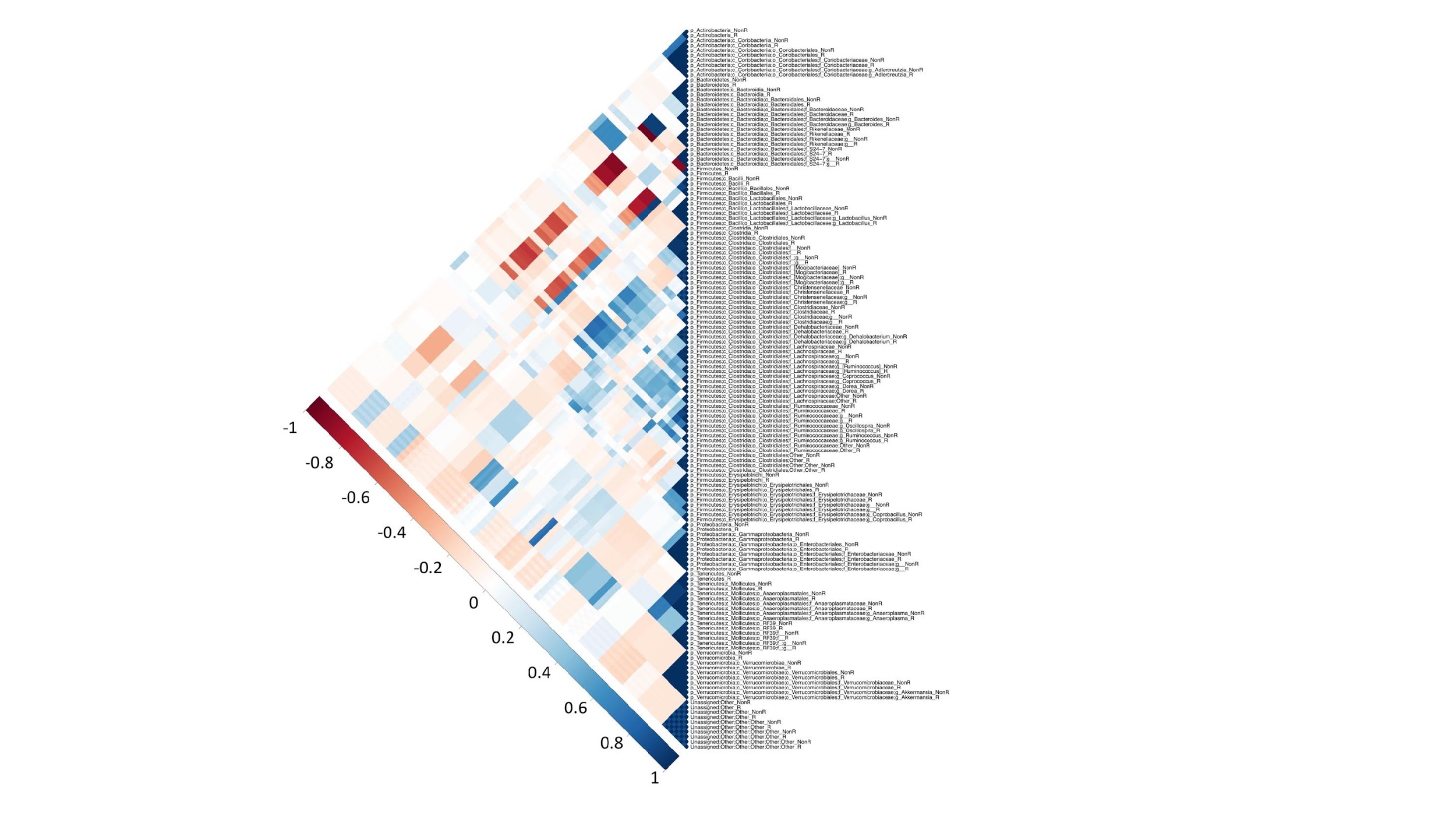

**Figure S3**. **Correlation plot between non-rarefied and rarefied taxa.** Heatmap depicting the correlations between non-rarefied (NonR) and rarefied (R) taxa show that the same taxa from both non-rarefied and rarefied datasets are strongly correlated, followed by taxa belonging to the same clade.
