## Supplementary material for "High-resolution QTL mapping with Diversity Outbred mice identifies genetic variants that impact gut microbiome composition": Fig S4

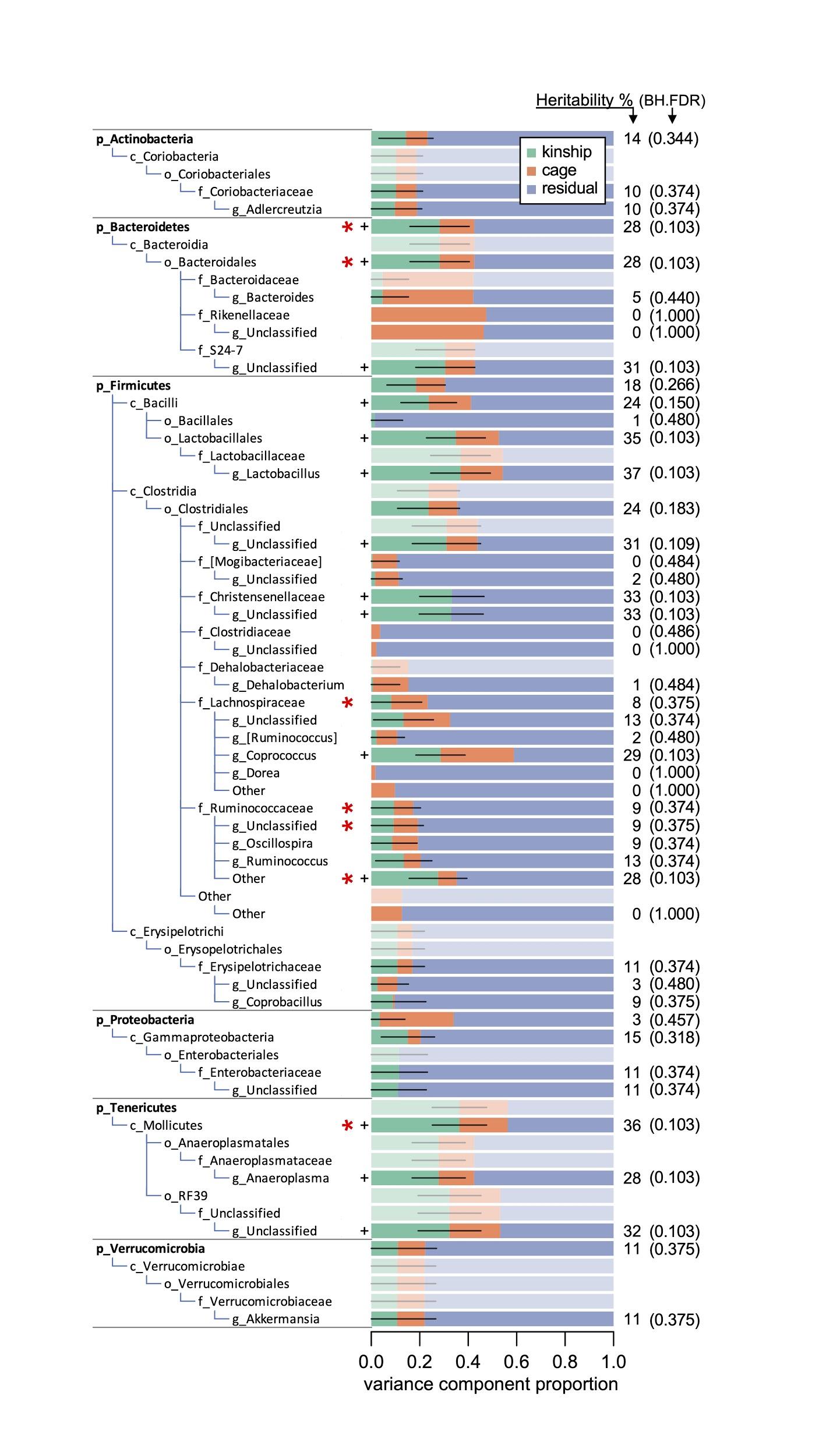

**Figure S4 - Proportion of variance estimates for kinship and cage for all taxa in rarefied data.** Proportion of variance estimates for kinship (green), cage effects (orange), and unexplained residual effects (blue) for each taxon. The kinship proportion of variance is an estimate of narrow sense heritability. Heritability percentages are shown on the left. Heritability standard error values are shown with black horizontal lines. Designations p_, c_, o_, f_, and g_ are for phylum, class, order, family, and genus, respectively. When results are identical across taxa in the same phylogenetic branch, only the lowest (most specific) taxa are shown and the rest are shaded out. Heritability significance is marked with one plus (**+**, RLTR *p*-value < 0.05) and BH FDR is shown in parentheses next to heritability percentages. Taxa marked with a red asterisk have statistically suggestive QTL (⭑, adj. *p*-value < 0.1). Complete table of heritability results, including non-rarefied data, can be found in **Table S3**.
