## Supplementary material for "High-resolution QTL mapping with Diversity Outbred mice identifies genetic variants that impact gut microbiome composition": Fig S5

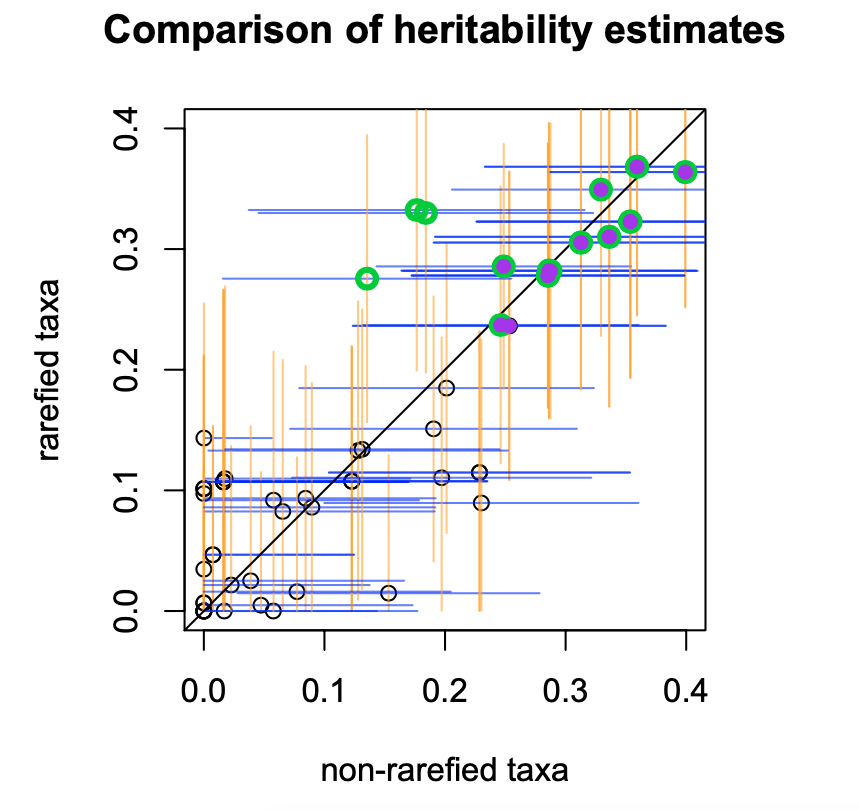

**Figure S5. Comparison of heritability estimates between non-rarefied and rarefied taxa**. Circles with purple fill correspond to non-rarefied taxa with statistically significant heritabilities. Circles with green outlines correspond to rarefied taxa with statistically significant heritabilities. Standard errors are shown as horizontal blue lines for non-rarefied taxa and vertical orange lines for rarefied taxa.
